# cfOncoXpress: Tumor gene expression prediction from cell-free DNA whole-genome sequences

**DOI:** 10.1101/2025.10.31.685915

**Authors:** Weiling Li, Marjorie Roskes, Alexander Martinez-Fundichely, Ekta Khurana

**Author notes:** These authors contributed equally to this work (Weiling Li, Marjorie Roskes, Alexander Martinez-Fundichely).

## Abstract

Cell-free DNA (cfDNA) fragments in the plasma capture cellular nucleosomal profiles since nucleosome-protected regions escape enzymatic degradation while nucleosome-depleted regions can not. We developed cfOncoXpress, a machine learning framework that uses fragmentation patterns to predict oncogene expression from cfDNA WGS. cfOncoXpress incorporates gene copy number aberrations inferred from cfDNA, including those associated with extrachromosomal DNA. Its application in prostate and breast cancers shows it can predict tumor subtype based on expression of signature genes and activated pathways. cfOncoXpress shows superior performance relative to other state-of-the-art methods and can be used to predict tumor gene expression when tissue biopsies are infeasible.

## Background

Cell-free DNA (cfDNA) refers to DNA fragments that circulate in the plasma and originate from cells undergoing apoptosis or necrosis during the maintenance of homeostasis^1,2^. In the context of cancer, cfDNA consists of both normal DNA and tumor-derived DNA (circulating tumor DNA, ctDNA)^3,4^. The study of cfDNA has become increasingly popular and has wide-ranging applications, such as cancer detection, tumor burden estimation, and disease progression monitoring. cfDNA is especially informative in the case of metastatic disease because of the elevated ctDNA content. Furthermore, its inherent capture of genomic and epigenomic information, coupled with its minimally invasive manner of collection allows overcoming the challenge of obtaining tissue biopsies from patients with late-stage disease when such procedures are not clinically feasible^5–10^.

cfDNA is enriched in genomic regions protected by nucleosomes, resulting in fragments of approximately ∼ 166-167 base pairs (bp)^11,12^. In closed chromatin, nucleosomes are typically spaced about 190 bp apart (length of nucleosomal + linker DNA)^12^; thus, cfDNA fragment depth at this periodicity can reflect the chromatin state of a given genomic region. In actively transcribed genes, promoter regions exhibit a nucleosome-depleted region (NDR) roughly 150 bp upstream of the transcription start site (TSS) with reduced nucleosome occupancy extending up to 1 kilobase into the gene body^13–15^.

Multiple studies have leveraged the unique fragmentation patterns of cfDNA towards a predictive end. In particular, two approaches have been developed to infer gene expression using cfDNA features. Ulz *et al*. employed a support vector machine (SVM) to classify genes as expressed or unexpressed based on cfDNA read depth coverage patterns at TSSs^16^. Because this method only gives binary classification to genes as expressed or unexpressed, it cannot be used to rank individual genes by expression level, making it difficult to accurately identify differentially expressed genes within or between samples. To overcome this limitation, Esfahani *et al*. developed EPIC-seq to infer individual gene expression from two features related to coverage and fragment length patterns at the TSSs^17^. EPIC-seq, a generalized linear model, was trained using a deep WGS cfDNA (200X) from one healthy control sample, and gene expression from a reference derived from a panel of peripheral blood mononuclear cells (PBMCs) from three different healthy controls. Before making a prediction, genes are grouped into sets of 10 based on their rank in the reference to improve performance. Thus, an expression value can only be predicted for each group of 10 genes. Grouping based on the control-derived reference, however, may lead to incorrect expression level predictions for upregulated oncogenes or downregulated tumor-suppressor genes in cfDNA samples from patients with cancer. Moreover, EPIC-seq uses targeted sequencing of genes of interest to study pathway activity, which requires prior knowledge of cancer-related information, which is often unavailable. While each approach has its own advantages and limitations, none of the models used >2 features, and matched cfDNA WGS and RNA-seq from tissue/cells of same individual for training. Importantly, although copy number (CN) has been shown to drive significant variability in gene expression^18–20^, neither of these approaches account for it in predicting gene expression. Tumors exhibit widespread copy number aberrations (CNAs) that are a major contributor to gene expression changes. Indeed, some of the extreme gene expression observed in oncogenes that drive tumor heterogeneity and therapeutic resistance is fueled by focal highly amplified complex DNA loci named amplicons, especially extrachromosomal DNA amplicons (ecDNA)^21–23^. ecDNA follows non-chromosomal inheritance and random segregation during cell division which promotes exceptionally high CN of oncogenes and their regulatory elements^23–25^. None of the existing methods incorporate ecDNA related CNAs inferred from cfDNA in gene expression prediction.

Moreover, other approaches to make accessibility-based inferences from cfDNA, such as Keraon^26^, rely on aggregating cfDNA signals across predefined genomic regions or regulatory sites, which cannot achieve site-specific resolution. Also, these models require reference profiles derived from patient-derived xenograft (PDX) models of the cancer type of interest, limiting their generalizability to new tumor contexts or samples lacking such reference data.

To overcome these limitations, in this study, we present a new method, cfOncoXpress, that incorporates both the fragmentation features of nucleosomal occupancy and CN to predict individual gene expression based solely on WGS cfDNA. cfOncoXpress applies a support vector regression model trained on deep WGS cfDNA from a healthy control sample^17^, followed by CN correction across samples. The use of healthy control samples, where the vast majority of cfDNA is contributed by PBMCs, is preferable for model training than the use of cfDNA from cancer patients due to the difficulty in estimating the contribution of ctDNA versus cfDNA from PBMCs in patient samples. While De Sarkar *et al*. employed PDXs where removing mouse reads from WGS results in nearly all reads from ctDNA^26^, we found that training on matched tumor tissue RNA-seq and cfDNA WGS obtained from a PDX did not yield optimal results for our approach, though this strategy enriches ctDNA content. Moreover, we demonstrate the utility of cfOncoXpress for subtype annotation of individual prostate cancer samples using a list of signature genes, and for pathway enrichment analysis for large cohorts of prostate (n=63) and breast (n=26) cancer cfDNA patient samples.

## Results

### Overview of patient samples used in the study

We used cfDNA from two cohorts of patients in this study: castration-resistant prostate cancer (CRPC) (n=63 plasma cfDNA samples)^27^, and ER+/HER2-metastatic breast cancer (MBC) (n = 26)^28^. The depth of sequencing for CRPC patients varies from 17.18X—182.39X (mean=106.01X, median=107.32X) and tumor fraction (TFx), estimated using ichorCNA^29^ based on copy number changes, ranges from 0.112—0.843 (mean=0.453, median=0.442) (Table S1). Additionally, 13 of these samples (sequencing depth: 72.11X—159.17X with mean=111.12X and median=111.61X, TFx: 0.112—0.635 with mean=0.396 and median=0.412) had matched tissue RNA-seq^27^ (Table S1). The CRPC cohort was classified into 33 adenocarcinoma (largely dependent on androgen receptor, AR) and 2 neuroendocrine (NE) patients based on histology. For the MBC patients, the depth of WGS varied from 2.46X—10.52X (mean=4.93X, median=4.55X) and TFx from 0.009—0.773 (mean=0.165, median=0.053)^28^ (Table S1).

### Multiple cfDNA features correlate with gene expression

#### Coverage around TSS

Nucleosomal depletion plays a crucial role in the dynamic regulation of chromatin structure and gene expression. Similar to Ulz *et al*., we define two regions of nucleosome occupancy reduction in the 2Kb region centered at the TSSs (*TSS* ± 1000 *bp*) called “2K-TSS” and a 200 bp regions (*TSS* − 150 *bp, TSS* + 50 *bp*) called “NDR” at which coverage can be used to distinguish between expressed and unexpressed genes^16^. Coverage within these regions was normalized by coverage per million bp to account for large regions with CN variation. We observe a depletion in coverage in the 2K-TSS region that is proportional to gene expression, and a dip in coverage immediately to the left of the TSS corresponding to the NDR (Fig. 1a and Fig. S1a).

**Figure 1.**
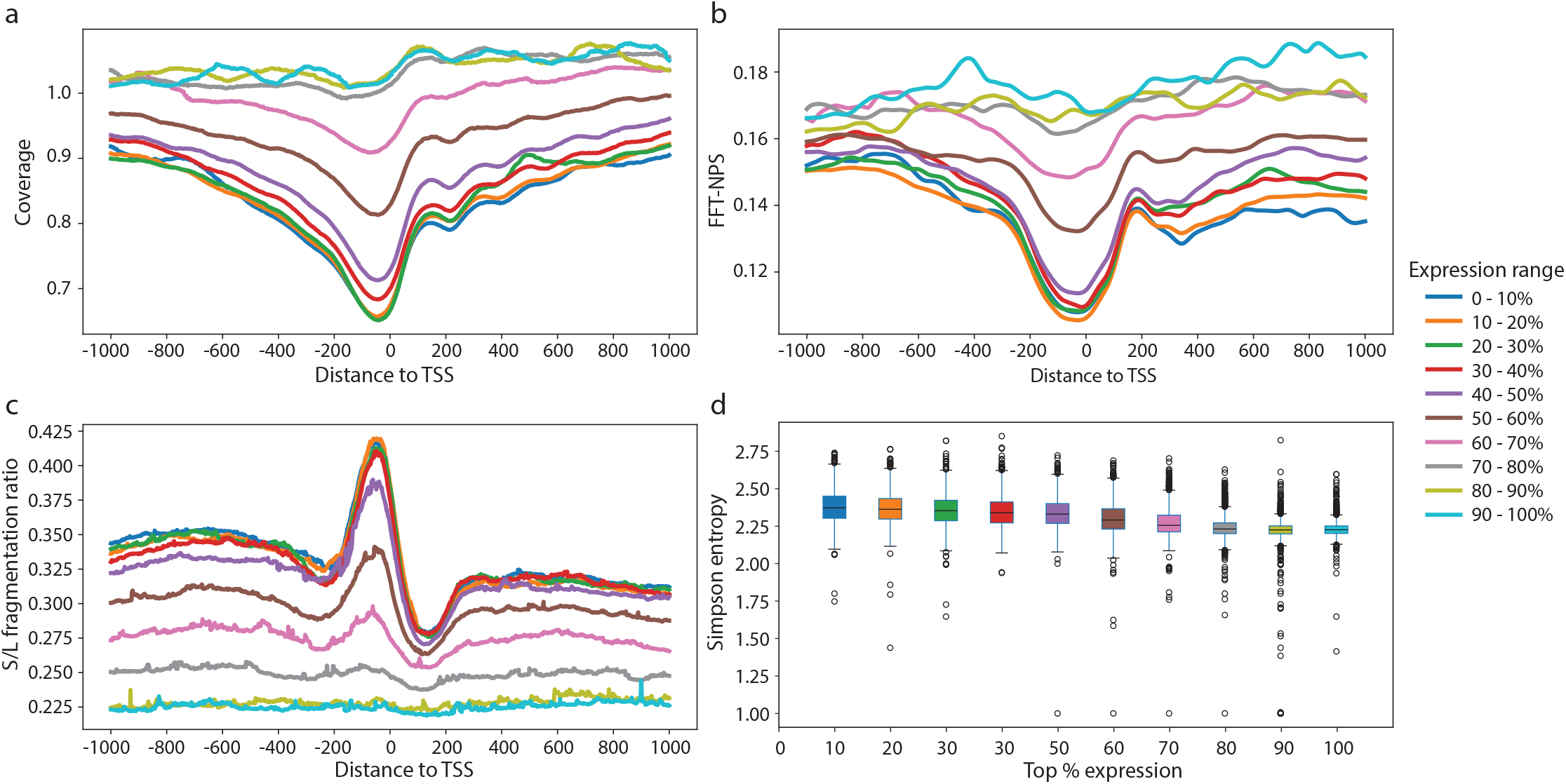
cfDNA features of a healthy example (control01) near transcription start sites correlate with gene expression. The list of expression percentiles follows the same color scheme in all panels. Fig. 1. a-c: the mean value of each feature (coverage, FFT-NPS, S/L) per expression decile at each base pair within the region (−1000 bp, +1000 bp) centered at TSS. d: boxplot of Simpson entropy of fragmentation according to different expression percentile. A higher Simpson entropy value indicates deviation from the dominance of the expected intermediate bin of nucleosome protection, which corresponds to higher gene expression.

#### Fast Fourier transformation (FFT) of the nucleosomal protection signal around TSS

The Window Protection Score (WPS), first described by Snyder *et al*, assigns a score to a window of the genome, quantifying its nucleosomal protection^12^. Because a window equal to the nucleosome length would exclude fragments misaligned by even a single base pair, we used a slightly smaller window (k = 120 bp) to allow minor positional variability of nucleosome-protected fragments. The WPS is defined as the difference between the number of fragments that span the entire window and the number of fragments with at least one endpoint in the window. The raw WPS values are post-processed to remove noise that arises from genomic events at larger and smaller scales^12^.To locate the regions of maximal nucleosome protection, the Nucleosome Protection Score (NPS) is defined as the smoothed WPS signal. Because nucleosomes have a 190 bp spacing in closed chromatin, we estimate the periodicity of nucleosome spacing by the amplitude of the FFT of the NPS, normalized by coverage per million bp at period 190 bp. We denote “2K-TSS FFT-NPS” as the FFT of the NPS for a 2 Kb window centered at TSS (*TSS* ± 1000 *bp*), and “NDR FFT-NPS” as the FFT of the NPS at a 3-nucleosome window encompassing the NDR (*TSS* − 380 *bp, TSS* + 190 *bp*). The FFT amplitude, not used by previous methods, indeed shows a lower value in the “2K-TSS FFT-NPS” region proportional to gene expression. As expected, FFT amplitude is lower in highly expressed genes corresponding to displaced nucleosomes at actively transcribed promoters (Fig. 1b and Fig. S1b).

#### Short-to-long (S/L) fragmentation ratio

This fragmentation metric reflects alterations in epigenetic and chromatin structure within transcriptionally active regions, in both normal and cancerous cells. As such, we used the short-to-long fragmentation ratio first described by Cristiano *et al*^30^. Short fragments are defined as those between 100 bp and 150 bp in length, whereas long fragments range from 151 bp to 220 bp^30^. This feature, denoted S/L fragmentation ratio, is positively correlated with gene expression levels (Fig. 1c and Fig. S1c).

#### Simpson entropy of fragment length

It has been observed that highly expressed genes show higher diversity in fragment length at their TSSs likely due to nucleosomal displacement and dynamics^17^. While Shannon entropy quantifies overall fragment size diversity, it lacks direct biological interpretability regarding degradation-associated fragment length. In contrast, Simpson entropy captures the diversity of fragment lengths associated with DNA degradation by accounting for their relative abundances within specific ranges of interest and reflects the degree of nucleosomal occupancy. We defined three bins of fragment-lengths: 100–170 bp reflects higher levels of DNA degradation, 170-210 bp reflects the expected fragment size associated with nucleosome protection and typical degradation levels, and 210–300 bp reflects reduced degradation. We formulate this metric as the inverse Simpson index: 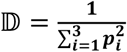, where *p*_*i*_ denotes the ratio of fragment length counts in bin *i* relative to total number of fragments across all bins. Since the intermediate bin represents nucleosomal protection, we expect a stronger dominance of this bin in unexpressed genes, yielding smaller entropy values at these promoters. Conversely, shifts toward higher or lower degradation weaken this dominance, resulting in higher Simpson entropy values. These changes reflect promoter activity and can serve as indicators of gene expression (Fig. 1d and Fig. S1d).

### cfOncoXpress: A support vector regression model to predict tumor gene expression from cfDNA WGS features

#### Integration of features using support vector regression

Using only WGS cfDNA, cfOncoXpress extracts the following features from promoter regions: 2K-TSS coverage, NDR coverage, 2K-TSS FFT-NPS, NDR FFT-NPS, S/L fragment ratio, and Simpson entropy of fragment length. These features are then used as input to machine learning models that predict individual gene expression in units of log_2_(TPM + 1) (TPM, transcripts per million). The model for predicting gene expression from cfDNA is ideally trained using matched samples where RNA-seq of all the cells that contribute cfDNA can be estimated. We achieved this using a deep WGS cfDNA sample (192.03X) from a healthy control with RNA-seq from matched PBMCs from the same healthy control^17^. Because the cfDNA signal from a healthy control is not confounded by heterogenous gene expression introduced by tumor-derived DNA, the model learns an unambiguous relationship between fragmentation patterns and gene expression. Five commonly used regression algorithms were implemented with hyperparameter tuning conducted via 5-fold cross-validation using Optuna^31^: gradient boosting regression (GBR), extreme gradient boosting (XGB), random forest (RF), support vector machine regression (SVR), and a neural network (NN) (Fig. 2). The accuracy of the predictions from each algorithm was evaluated by the Spearman correlation and mean squared error (MSE) between the gene expression predictions and the observed expression from matched PBMC RNA-seq. Although NN achieved the highest Spearman correlation and lowest MSE between predicted and observed gene expression on the internal validation set (20% of control01 regions; Fig. S2), SVR achieved the highest Spearman correlation in patient samples, while maintaining an acceptable MSE (Fig. S3a,b). This indicates that other models exhibited signs of overfitting, and SVR was therefore used for all further analyses.

**Figure 2.**
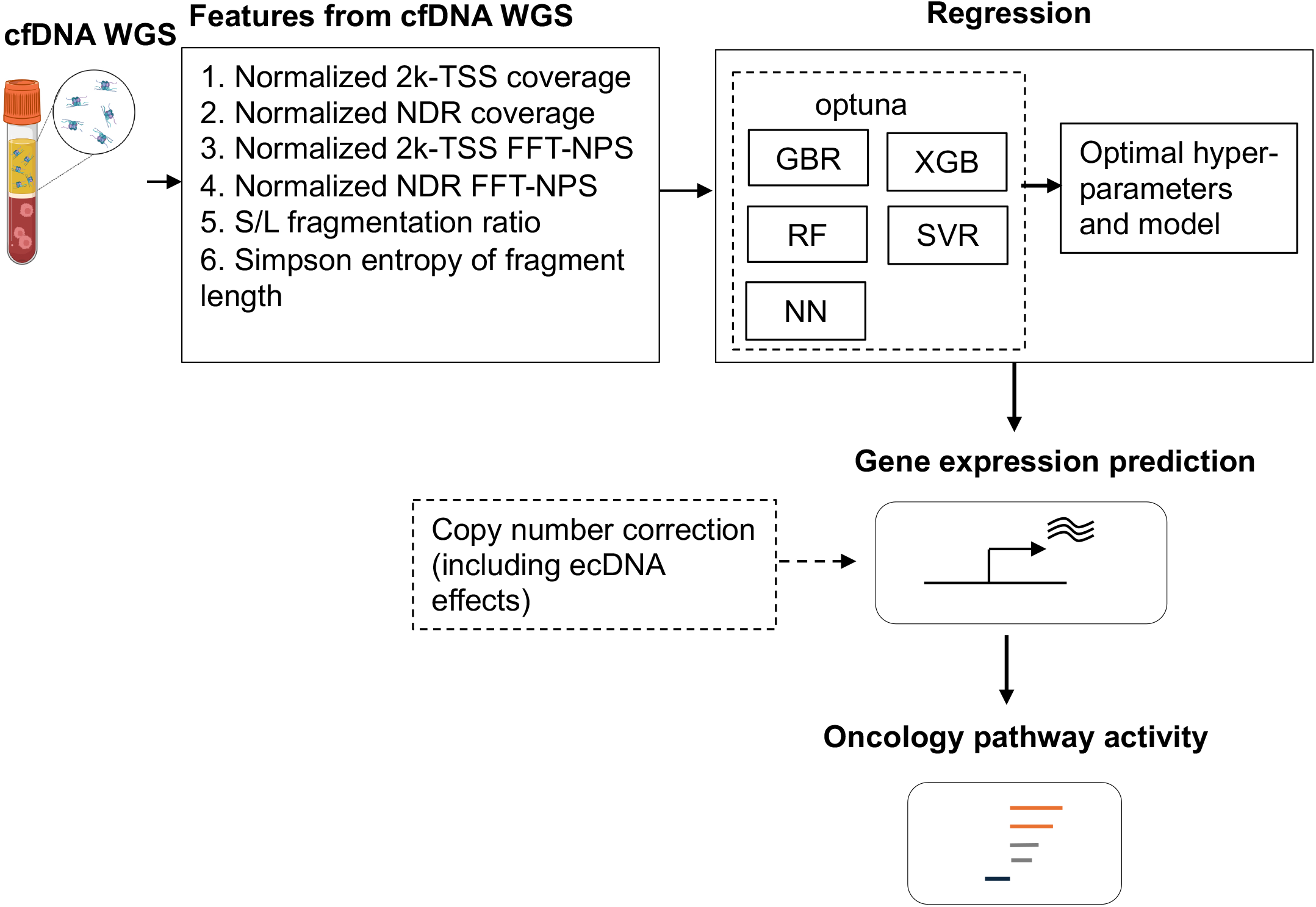
Overview of the cfOncoXpress pipeline. cfOncoXpress takes only cfDNA WGS as input, from which six promoter-associated cfDNA features are extracted. These features are used to train regression models (i.e., Gradient Boosting regression (GBR), Extreme Gradient Boosting (XGB), Random Forest (RF), Supervector Machine Regression (SVR) and a Neural Network (NN)), with hyperparameter optimization performed using Optuna to identify the best-performing model and parameter set for gene expression prediction. A copy number correction is applied post-regression to further improve accuracy. Effects of ecDNA can also be evaluated in downstream analysis. Finally, the predicted gene expression values can be used to infer pathway activity.

Another approach to train the model can be the use of cfDNA WGS sample from PDXs with matched tumor tissue RNA-seq since the removal of mouse reads from WGS results in nearly all reads from ctDNA^26^. We find that while a model trained using healthy control and prostate cancer PDXs shows a modest improvement in performance with regard to Spearman correlation and MSE between predicted and observed gene expression (Fig. S4), it does not generalize as well to other cancer types (see Methods). Thus, cfOncoXpress uses the model trained using only the healthy control sample.

#### Copy number correction and ecDNA effects

It is well known that copy number (CN) alterations can often lead to significant variability in the expression of genes they impact. It is important to note that this effect is particularly relevant for oncogenes which are often amplified during cancer progression, and this is especially the case in the context of extrachromosomal DNA (ecDNA) amplicons^41^. While we observe no correlation (Pearson) between gene CN and its expression across genes in a given sample (Fig. S5), we do observe correlation (Pearson) between CN and expression for a given gene across samples (Fig. 3a, Table S2). Therefore, CN correction was applied only after the initial SVR-based predictions were made.

**Figure 3.**
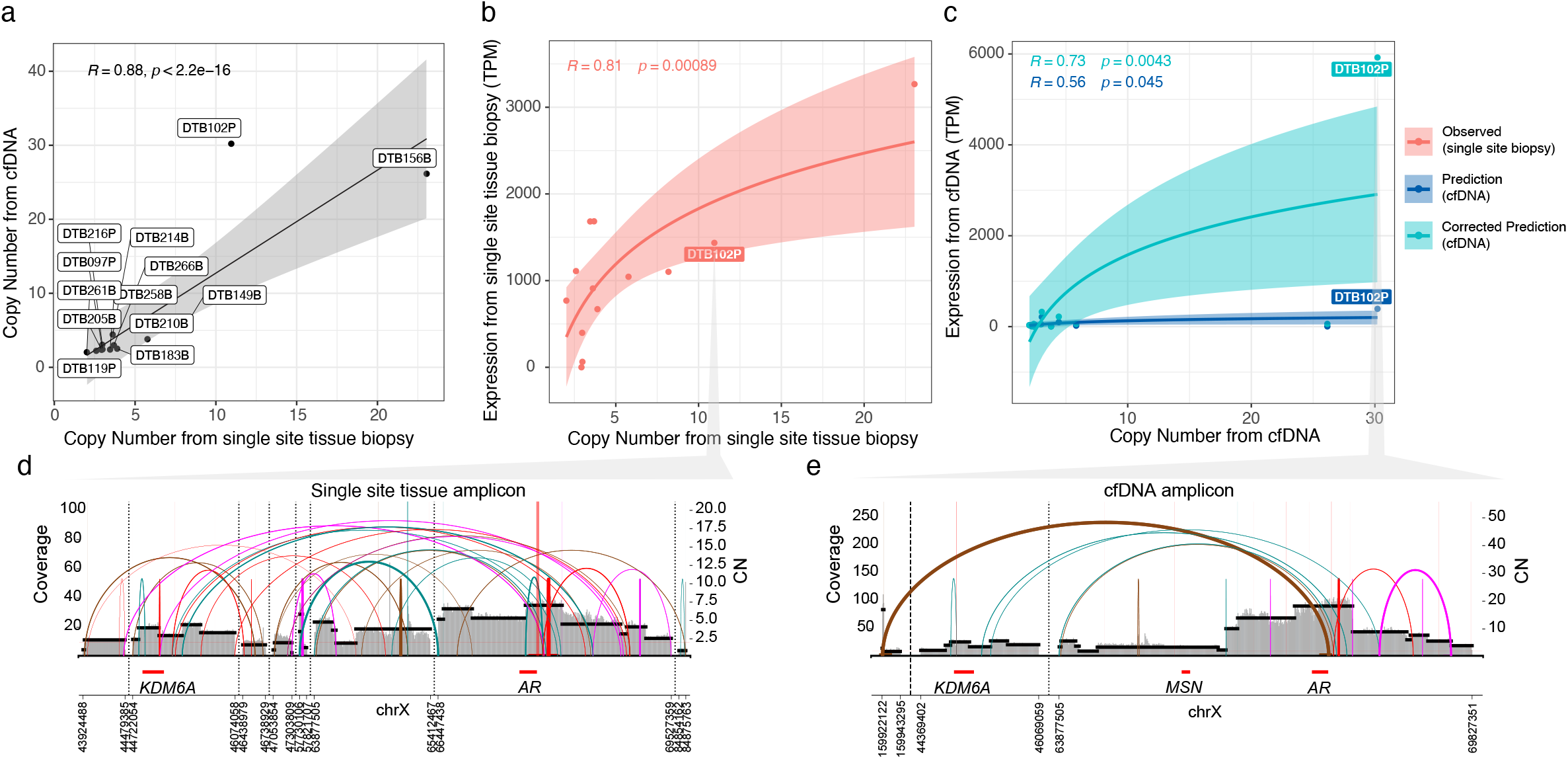
AR Copy number correction and ecDNA effects. a Pearson correlation between the copy-number derived from cfDNA and tissue biopsy. b Impact on AR expression of the corresponding copy number for each sample. c Predicted AR expression from cfDNA and copy number correction for these values. d ecDNA amplicon affecting AR detected in the tissue biopsy of sample DTB102P. e ecDNA amplicon affecting AR detected from the cfDNA of sample DTB102P.

For each cfDNA WGS sample, and matched WGS tissue sample when available, we estimated CN alterations using CNVkit^32^ and ecDNA amplicons using the AmpliconArchitect Suite^33^. 92% (n = 12) of samples with matched WGS cfDNA and tissue RNA-seq showed amplicons in at least one of the data types (Fig. S6a). These results for tissue WGS are consistent with those previously reported on the same cohort of prostate cancer patients^34^. However, to our knowledge, our study is the first to demonstrate the presence of ecDNA in tumor cells using cfDNA WGS. We detected amplicons in eight cfDNA samples, five of which had corresponding amplicons in matched tissue (Fig. S6a). We hypothesize that the discordance in the number and structure of amplicons detected from different data sources may arise from both biological reasons, and technical differences in assembly confidence when reconstructing complex ecDNA and linear amplicon structures from cfDNA WGS versus tissue WGS (Fig. S6b). A biological plausibility is that cfDNA provides a comprehensive sampling of genomic alterations across multiple tumor sites within the metastatic landscape, whereas single-site tissue biopsies are spatially constrained and may not capture the full spectrum of amplicon heterogeneity present across anatomically distinct lesions. For technical differences, the inherent fragmentation patterns and coverage heterogeneity in cfDNA may impact the ability to call structurally complex amplicons compared to the more uniform coverage depth achieved in tissue WGS. Importantly, we observe strong correlation between CN from cfDNA WGS and matched tissue WGS (Pearson correlation 0.803—0.954, Fig. 3a, Fig. S7 and Table S3), confirming that cfDNA can capture CN changes occurring in the tumor tissue. For example, the *AR* gene, a key CRPC oncogene, showed markedly higher gene expression in tissue in samples with *AR* amplifications (Fig. 3b). In particular, we highlight sample DTB102P as an example where similar ecDNA amplicons were identified from both cfDNA and tumor tissue WGS^27^ (Fig. 3d,e). Notably, the *AR* oncogene was found within these ecDNA, leading to a substantial increase in its copy number.

Among all 63 CRPC cfDNA samples, ecDNA was detected in 10, 7 of which had ecDNA amplicons at the *AR* locus. An additional 20 more samples exhibited other types of amplicons at the *AR* locus, such as linear amplicons that are highly amplified focal DNA segments^35–37^, breakage fusion bridge (BFB) amplicons originated during repetitive cycles of breakage and fusion^35–38^, and complex-non-cyclic amplicons which are highly amplified complex DNA structures^35–37^ (Table S4).

To account for CN alterations, we multiply the gene expression predictions from SVR by their corresponding CN values:

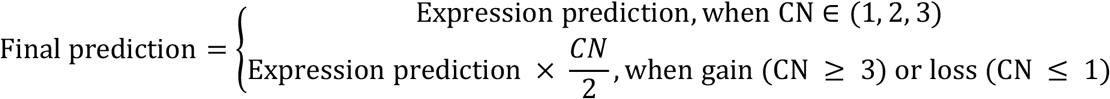

This correction effectively normalizes for gene dosage, improving the concordance between predicted and observed expression, thus improving the overall accuracy of the method (Fig. 3c, Table S5). The effect of this correction is particularly stark in a number of genes implicated in prostate cancer (Fig. S6). These include *MYC*, a major driver of tumorigenesis and progression^39^; *YIPF6*, which has been reported to be co-amplified with the AR-V7 splice variant^40^; *EFNB1*, whose upregulation is associated with increased cell migration and invasion^41^; and *OPHN1*, whose overexpression promotes development of CRPC and can be induced by androgen deprivation therapy (ADT)^42^. More examples include *FAM155B*, whose overexpression is associated with poor prognosis^43^; *USP11*, which promotes progression by up-regulating *AR* and *MYC* activity^44^; *NDUFB11*, which has been identified as a potential driver gene in aggressive disease^45^; and *KDM6A*, which plays a crucial role in the maintenance of *AR* activity^46^ (Fig. S6).

By comparing gene expression predictions in samples from patients with cancer to the RNA-seq gene expression of a panel of PBMCs from healthy controls, we can perform pathway enrichment analysis. We present an overview of the cfOncoXpress pipeline in Fig. 2.

### Application and comparison of cfOncoXpress predictions with those from other methods

The availability of matched cfDNA WGS and tissue RNA-seq from patient samples enables evaluation of cfOncoXpress gene expression prediction in patients with cancer^27^. These data provide an excellent resource for model evaluation and comparison with other state-of-the-art methods.

#### A. Correlation of expression predictions from cfDNA WGS with RNA-seq from matched tumor tissues of same patients

To evaluate the validity of cfDNA-derived predictions, we compared them to the gene expression profiles from RNA-seq of matched tumor tissues from 13 samples from the CRPC cohort^27^, using Spearman correlation coefficient for our non-linear model and MSE as metrics of performance (Fig. 4a,b). Ten of thirteen samples show strong correlation between predicted and expected gene expression (Spearman R = 0.5 – 0.6), while two samples show only moderate correlation (Spearman R < 0.45). Importantly, all samples do better than the random expectation of no correlation (Fig. 4a).

**Figure 4.**
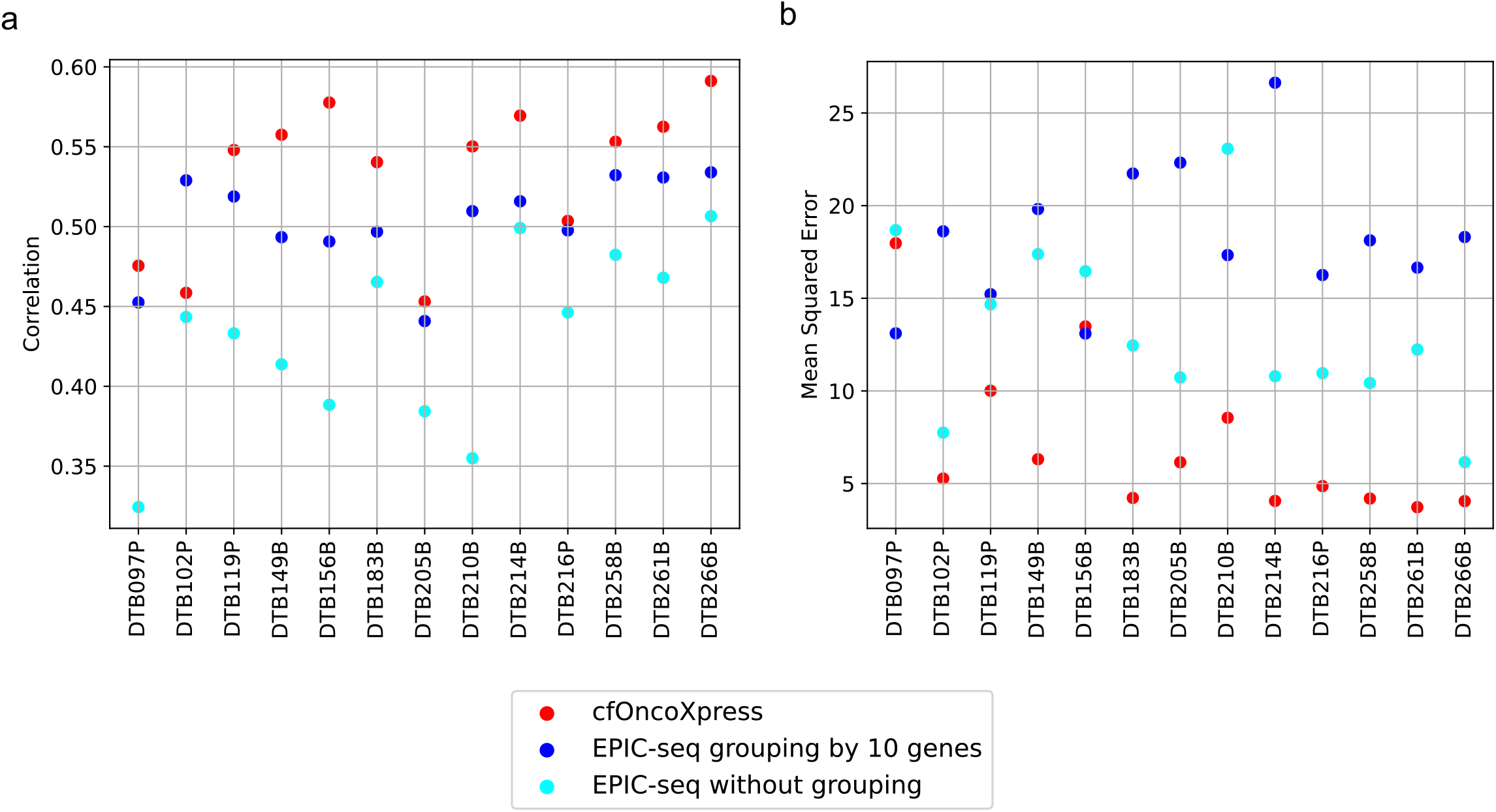
Model performance was evaluated using Spearman correlation and mean squared error between prediction and log_2_(TPM+1) . (a) (b) Sample ids are from Herberts *et al*^2^. P represents “Progression”, and B represents “Baseline”, related with the tumor development. Generally speaking, cfOncoXpress can achieve higher correlation while lower MSE.

Next, we benchmarked the performance of cfOncoXpress against EPIC-seq^17^, the only previously reported method for inferring expression of individual genes from cfDNA. The method by Ulz et al.^16^ classifies genes as highly expressed or unexpressed and cannot be used to compute correlation of predicted versus observed expression values. We observe higher correlation between predicted and expected gene expression in twelve of thirteen samples using cfOncoXpress compared to EPIC-seq (Fig. 4a). Furthermore, eleven samples show lower MSE using cfOncoXpress compared to EPIC-seq (Fig. 4b). Importantly, while much of the correlation achieved by EPIC-seq relies on its grouping by 10 genes, this grouping leads to higher MSE compared to non-grouped genes in the majority of samples (10/13) (Fig. 4a,b).

#### B. Detection of subtype-specific signature genes

To assess the ability of cfOncoXpress to capture biologically meaningful distinctions between disease subtypes, we compared the predicted and observed gene expressions of subtype-specific genes in these 13 CRPC samples none of which show NE histology^27^. We considered 93 CRPC-AR and 93 CRPC-NE signature genes^47^. To establish a baseline of expectation, we first compared these gene sets in the tissue RNA-seq data, and observed significantly higher CRPC-AR signature gene expression than CRPC-NE signature gene expression in 77% (10 of 13) of samples (one-sided Mann–Whitney U test, Fig. 5). Then, by comparing the cfOncoXpress predictions for these gene sets, we observe 7/10 samples with significantly higher CRPC-AR signature gene expression than CRPC-NE (Fig. 5). Of the three samples that did not achieve statistical significance, two (i.e. DTB258B and DTB266B) had low tumor fraction (< 0.2) and the other (i.e. DTB183B) may reflect mixed plasma signals from multiple tumor sites of different subtypes (unlike RNA-seq which is derived from a single-site tissue biopsy).

**Figure 5.**
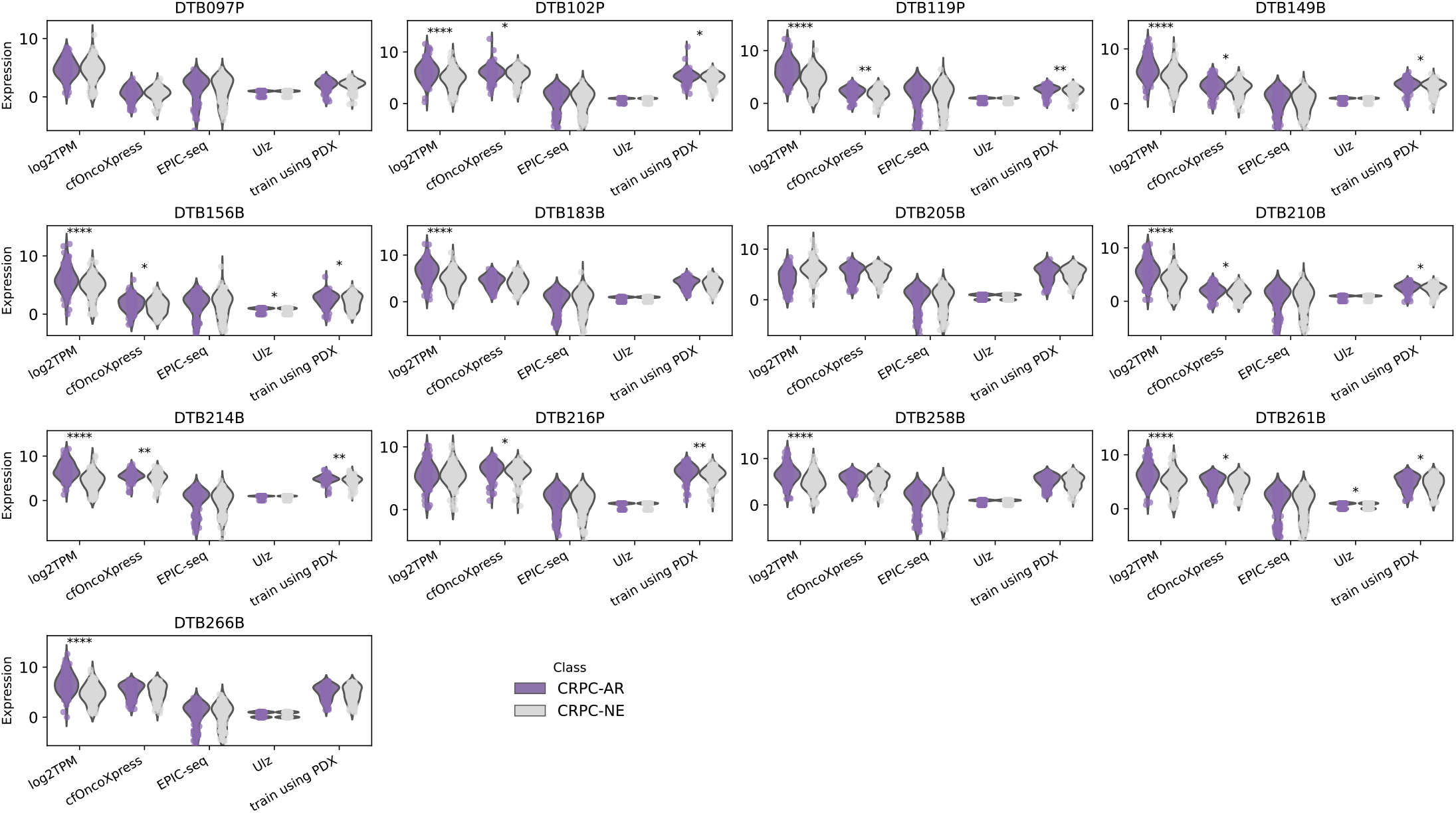
A one-sided Mann–Whitney U test was applied to assessing whether CRPC-AR expression is significantly higher than CRPC-NE. *: 0.01 < p <= 0.05; **: 0.001 < p <= 0.01; ***: 0.0001 < p <= 0.001; ****: p <= 0.0001. (a) In RNA-seq data, 10 out of 13 adenocarcinoma patients showed significantly higher CRPC-AR expression compared to CRPC-NE, with exceptions in DTB097P, DTB205B, and DTB216P. (b) Among these 10 patients, cfOncoXpress predictions identified 7 with significantly higher CRPC-AR expression. DTB258B and DTB266B did not show significant differences, likely due to low tumor fractions (<0.2). (c) EPIC-seq predictions did not show significant differences in any sample. (d) Using Ulz’s classification method, most signature genes were predicted as expressed (1) for either CRPC-AR or CRPC-NE, resulting in only two samples (DT156B and DTB261B) showing significant differences. (e) When trained on controls and LuCaP PDXs, the results are same as from cfOncoXpress.

For comparison with other methods, we also compared expression of these signature gene sets using predictions from EPIC-seq^48^ and the Ulz *et al*^49^. EPIC-seq predictions were not significantly different in the CRPC-AR and CRPC-NE gene sets in any of the samples (Fig. 5). Ulz *et al*.’s binary expression predictions (achieved for 51–78% of genes) were only significantly different in the CRPC-AR and CRPC-NE gene sets in two samples (Fig. 5). Overall, cfOncoXpress performs better than both these other methods to distinguish subtype-specific expression patterns.

### Impact of tumor purity and sequencing depth on prediction accuracy

To estimate the lower bounds of sequencing depth and tumor fraction for reliable gene expression prediction, we performed *in silico* downsampling experiments using a patient sample (DTB214B, 125.3X, TFx = 0.63) (Methods). Performance was considered robust when the MSE was lower than 4.404, a threshold derived from the SVR’s best performance (Methods, Fig. S2d). Using these criteria, we observe good performance for sequencing depth as low as 25.066X and tumor fraction down to 0.20 (Fig. 6).

**Figure 6.**
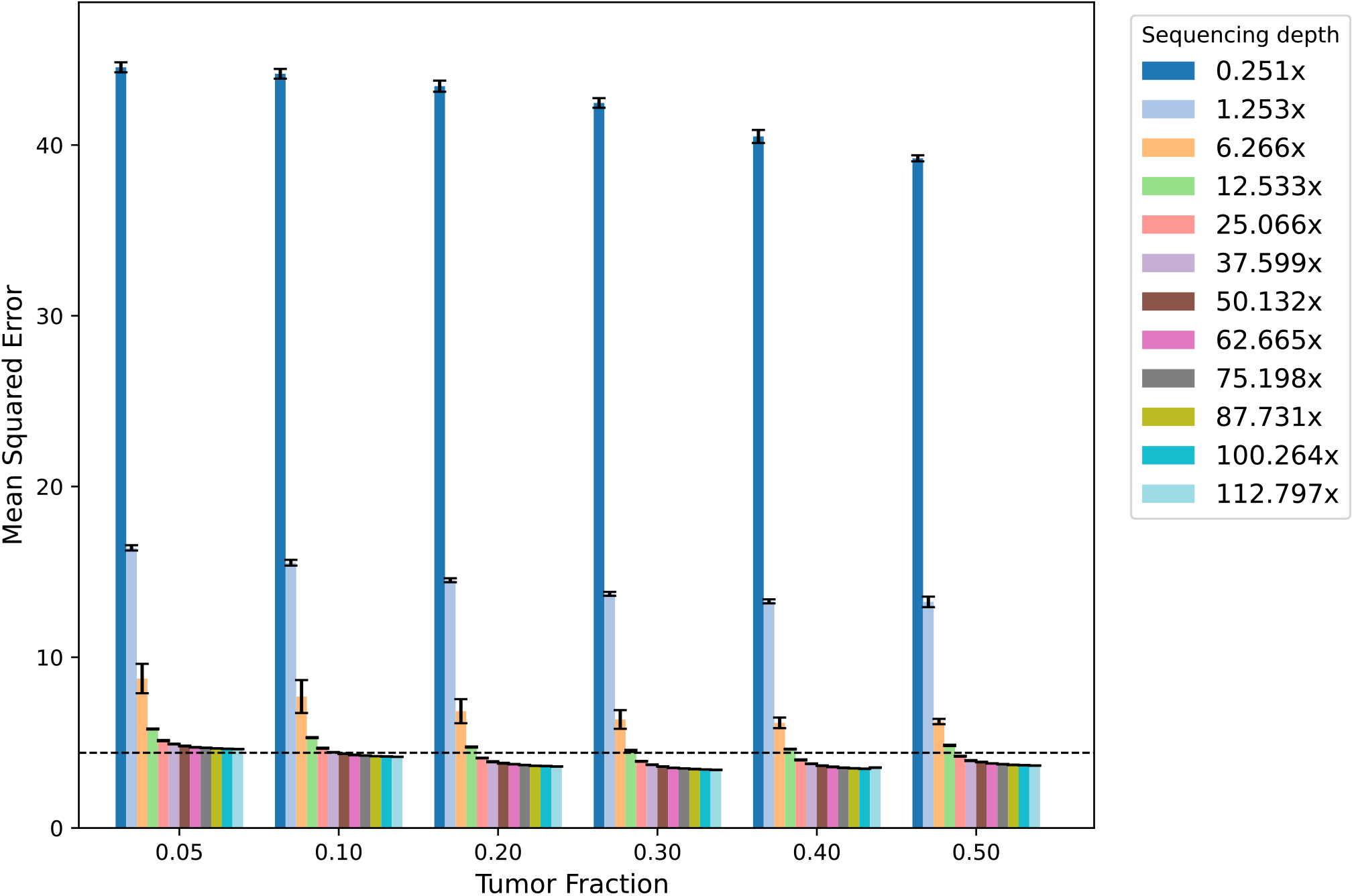
Mean squared errors (MSE) for different sequencing depth and tumor fraction in downsampled DTB214B. For each combination of sequencing depth and tumor fraction, we generated 5 synthetic samples. Error bars are shown for each setting. The dashed line represents an MSE of 4.404, corresponding to the point where Support Vector Regression (SVR) achieved the best performance in the validation experiment (Fig. S2d). Data points below this line indicate better performance, with sequencing depth ≥ 25.066x and tumor fraction ≥ 0.2.

### Application of cfOncoXpress for pathway enrichment analysis

Having established that cfOncoXpress outperforms existing methods in predicting gene expression and detecting enrichment of subtype-specific genes, we next investigated if the predictions were robust enough for pathway activity analysis. This is especially relevant as we envision that one of the most common uses of cfOncoXpress predictions in new patient cohorts would be to infer dysregulated biological pathways. Thus, using Gene Set Enrichment Analysis (GSEA)^50^, we compared disease-specific pathway activity using the predicted gene expression in these cohorts as described above – deeply sequenced CRPC adenocarcinoma (CRPC-Adeno, n = 61) and neuroendocrine (CRPC-NE, n = 2) samples, and a shallowly sequenced ER+/HER2-MBC cohort (n = 26). We note that we used the published cohorts of cfDNA WGS from patients with metastatic cancer only since the tumor fraction may be too low for accurate predictions in patients with primary cancer only. To identify differentially expressed genes and enriched pathways, we compare our predicted expression in patients with the RNA-seq data of PBMCs from healthy participants. In healthy individuals, the vast majority of cfDNA is contributed by PBMCs, and thus comparison with this dataset points to pathways related to cancer. The panel includes RNA-seq data from PBMCs from 34 healthy participants^51–53^.

We observe upregulation of the TCGA Androgen Response pathway in the CRPC-Adeno cohort, which is consistent with AR driving the growth of these tumors (Fig. 7a). As expected, this pathway is not upregulated in either the CRPC-NE or MBC cohorts. Conversely, while we observe upregulation of the Beltran NEPC (neuroendocrine prostate cancer) Up pathway^54^ in the CRPC-NE cohort, we do not see this trend in the CRPC-Adeno or MBC cohorts (Fig. 7b). Similarly, we observe upregulation of the SMID Breast Cancer Luminal A Up pathway^55^ in the MBC cohort alone, which is consistent with the fact that luminal A tumors represent the dominant molecular subtype of ER+/HER2– breast cancers^56^ (Fig. 7c). Thus, we have demonstrated that cfOncoXpress may be used to infer pathway enrichment in metastatic cancer patients in the absence of tissue RNA-seq.

**Figure 7.**
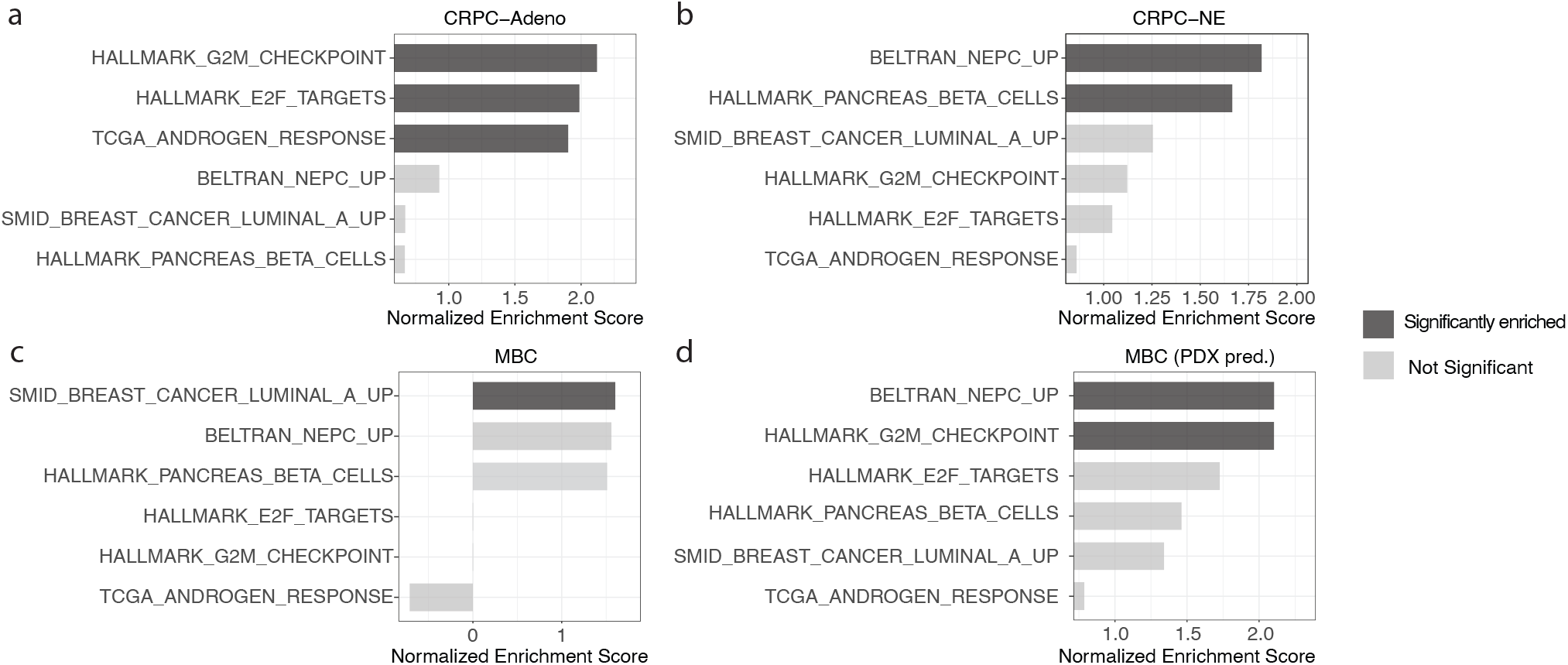
Pathway enrichment analysis using GSEA. The input datasets for pathway enrichment analysis are the predictions of prostate cancer (CRPC), and breast cancer (MBC) cohorts, and the RNA-seq expressions from a panel f healthy controls. Significant enrichment was defined as NOM p-value ≤ 0.05, FWER p-val ≤ 0.05, and FDR q-value ≤ 0.05. (a) Pathways enriched in CRPC-Adeno compared to the other cohorts when using cfOncoXpress, where TCGA_ANDROGEN_RESPONSE (CRPC-AR) is significantly positively enriched. (b) Pathways enriched in CRPC-NE relative to the others when using cfOncoXpress, where BELTRAN_NEPC_UP (CRPC-NE) is significantly positively enriched. (c) Pathways enriched in the ER+/HER2− MBC cohort compared to the rest when using cfOncoXpress, where SMID_BREAST_CANCER_LUMINAL_A_UP is significantly positively enriched. (d) A comparable pathway enrichment analysis was performed using predictions from a model trained on controls and LuCaP PDX data; however, this analysis failed to detect significant enrichment in MBC as BELTRAN_NEPC_UP (CRPC-NE) is positively enriched here.

### Number of patient samples needed for pathway enrichment analysis

Next, we performed power calculations to estimate the patient cohort size needed for pathway activity analysis given the panel of PBMC RNA-seq from healthy controls (n = 34) against which to test enrichment. This was done by modeling the expression distributions of tumor and healthy controls groups (Methods). On average, the predicted expression of genes in the CRPC-AR pathway (i.e., TCGA_ANDROGEN_RESPONSE) was 2.83-fold higher in the CRPC-Adeno cohort than in the panel of healthy controls, suggesting at least three patient samples are required to achieve statistical significance. In contrast, the predicted expression of genes in the CRPC-NE pathway (i.e.,Beltran_NEPC_UP) was 5.08-fold higher in the CRPC-NE cohort than in the panel of healthy controls, suggesting that just a single patient patient sample is required to achieve statistical significance for this CRPC subtype. Similarly, the predicted expression of genes in the SMID Breast Cancer Luminal A Up pathway was 3.35-fold higher in the MBC cohort than in the panel of healthy controls, suggesting only two patient samples are required to achieve statistical significance (Fig. 7d). These results highlight the robustness of cfOncoXpress in capturing subtype-specific pathway activity, even from small patient cohorts.

## Discussion

In this study, we present cfOncoXpress, a computational method that utilizes cfDNA WGS from minimally invasive liquid biopsies for gene expression prediction. cfOncoXpress incorporates six features obtained from cfDNA WGS into an SVR to predict individual gene expression. Because CN alterations play a critical role in gene expression by impacting the abundance of individual genes, they are incorporated in the expression predictions. This study is the first, to our knowledge, to present and incorporate ecDNA associated CN alterations from cfDNA WGS. We show the application of cfOncoXpress on metastatic prostate and breast cancer samples, and demonstrate its superior performance relative to other state-of-the-art approaches.

We have demonstrated several use-cases, including pathway activity analysis across samples as well as identification of subtype-specific gene activity within samples. cfOncoXpress will be extremely useful in monitoring disease progression as metastatic tumors often shed large amounts of DNA into the plasma, especially since, unlike tissue biopsies, serial cfDNA sample collection is eminently feasible. Using downsampling calculations, we show the ability of cfOncoXpress to predict gene expression across a wide range of sequencing depths and tumor fractions demonstrating the robustness of our approach.

We note that while gene CN estimated from cfDNA WGS is highly correlated with CN estimated from tissue WGS (Fig. S7, Table S3), there may be discordance between cfDNA-derived CN estimates and those obtained from single-site tissue (Fig. 3a). This may be the result of both technical and biological factors. From a technical perspective, inherent noise in the signal can make it more challenging to confidently identify CN alterations in cfDNA relative to equivalent analyses performed on WGS tissue, which is especially the case in samples with lower tumor fraction (Fig. S7). From a biological perspective, single-site tissue biopsies may yield underestimated copy number values in CRPC patients with multiple metastatic sites, where the inter-tumor heterogeneity may present variable CN alterations across different sites. In contrast, cfDNA has the potential to capture the full heterogeneity of the metastatic landscape.

Nonetheless, the signal from cfDNA is able to capture samples with gene amplifications even if the exact CN estimate may vary from the single-site tissue biopsy signal.

Genome-wide studies have identified NDRs at the TSSs of actively transcribed genes, which is the basis for using cfDNA fragmentation patterns at promoters for gene expression prediction. However, we expect the same phenomenon to be observed at distal enhancer regions where nucleosome depletion is accompanied by binding of transcription factors to modulate gene expression^57–61^. Currently, cfOncoXpress as well as previous related studies, such as EPIC-seq^17^, and Ulz *et al*.^16^, only utilize cfDNA features at the gene promoters. Looking ahead, we will incorporate cfDNA features at both promoters and enhancers in cfOncoXpress to further increase gene expression prediction accuracy.

## Conclusions

cfOncoXpress is a machine learning framework that uses cfDNA WGS to predict gene expression of cells undergoing apoptosis and contributing to cfDNA. By comparison with a panel of healthy controls, it can predict tumor-specific gene expression. It incorporates CN of oncogenes inferred from cfDNA WGS for expression prediction. We have shown that cfOncoXpress outperforms other existing methods for gene expression prediction. We present practical use cases of the predictions, such as identification of subtype-specific gene enrichment within a sample, as well as identification of activated pathways in a given patient cohort. cfOncoXpress can predict reliable expression values for sequencing depth as low as 25X and tumor purtiy as low as 0.20 demonstrating its utility in guiding therapeutic decisions.

## Methods

### Cohorts

#### Healthy control

For model training, we used deep cfDNA WGS from a healthy control (192.03X) together with matched PBMC RNA-seq from Esfahani *et al*.^17^ (Table S1) due to the difficulty in accurately estimating the relative contributions of ctDNA and PBMC-derived cfDNA in cancer samples.

#### Patient CRPC cohort

We applied cfOncoXpress to 63 cfDNA samples from 35 patients with CRPC reported by Herberts *et al*.^27^, on which they performed deep WGS (17.18X-182.39X, mean = 106.01X). Tumor fractions were estimated using ichorCNA^29^ (0.112-0.8428, mean = 0.453) (Table S1). Two of these samples were from patients with neuroendocrine histology^27^. Time-matched tissue RNA-seq for 13 samples were also provided^62^ (Table S1).

#### Patients MBC Cohort

Shallow WGS of cfDNA from 26 patients with ER+/HER2-MBC were obtained from Bujak *et al*. (NCBI BioProject accession: PRJNA578569)^28^ (Table S1).

#### LuCaP PDX

One LuCaP PDX from each subtype with a high tumor fraction was selected for evaluation using LuCaP PDX data in the training set. Specially, one CRPC-AR (LuCaP_92, 20.13X, TFx= 0.8287) and one CRPC-NE (LuCaP_208.4, 36.95X, TFx = 0.9694) PDX samples were obtained from De Sarkar *et al*.^*63*^ (Table S1).

#### Panel of healthy controls

To identify differentially expressed genes and enriched pathways, we compare cfOncoXpress predicted gene expression from patient cfDNA samples against the gene expression from RNA-seq of PBMCs from 34 healthy controls from various publications (GSE107011^51^, GSE115259^52^, PRJNA739257^53^) (Table S1).

### Optuna hyperparameter tuning

Hyperparameter tuning was performed using the Optuna^64^ Python package with a 5-fold cross-validation strategy applied to the training data. A total of 150 optimization trials were conducted on each of the five regression models (i.e., gradient boosting, extreme gradient boosting, random forest, support vector regression, and a feed-forward neural network using Tensorflow^65^) to simultaneously maximize the Spearman correlation and minimize the MSE using a multi-objective setup. Optimal hyperparameter values were selected based on the performance observed across these trials for each model (Fig. S2). For models tuned using multi-objective optimization with Optuna, where a Pareto front of optimal solutions may be returned, we prioritized the solution with the highest Spearman correlation to best preserve the rank order of gene expression. These top-performing hyperparameters were then applied to their respective regression models and evaluated on patient data. Among all models, SVR achieved the highest Spearman correlation between predicted and observed gene expression values, while maintaining an acceptable MSE, and was, therefore, selected as the best-performing model (Fig. S3a, b).

### Feature Importance

Permutation feature importance, which assesses the impact of each input feature by randomly shuffling values and measuring changes in model performance, was used to interpret the SVR model with a radial basis function (RBF) kernel. Since RBF is a non-linear and non-parametric model, traditional coefficient-based importance is not applicable. Features that, when permuted, cause a large drop in performance are considered more important. This approach provides a model-agnostic and intuitive approach to evaluating the contribution of individual input features to the predictive power of the SVR model. We observed that coverage-based features were the most important in accurately predicting gene expression (Fig. S3c).

### CN and DNA amplicon detection

CN detection was performed using CNVkit^32^ (v0.9.7), and amplicon detection was performed using AmpliconSuite^33^ (v1.3.2). We used the CN calls from CNVkit with a CN > 5 and 10 Kb < size < 10 Mb in which to search for amplicons. The result classifies the amplicons in four categories, including: ecDNA, linear, BFB, and complex-non-cyclic amplicons, explained in the “Copy number correction and ecDNA effects section” above.

### Implementation and parameter configuration of other methods for expression prediction from cfDNA (EPIC-Seq and Ulz)

The parameter settings were consistent with those employed in the original EPIC-Seq study.

Rscript runEPIC.R --bamdir $bamdir/ --tssinfo priordata/all.tss.genes.canonical.ensembl75.txt -- panelbed selectors_windows/all.tss.genes.canonical.ensembl75.selector.txt --targeted no –outdir $outdir/ --mapq 30 --groupref pbmc_genome --groupsize $groupsize ($groupize = 1 or 10)

Rscript runGEPModel.R --xnewpath $outdir/epicseq.features.merged.by.$groupsize.txt --tssinfo priordata/all.tss.genes.canonical.ensembl75.txt --outdir $outdir/

We applied Ulz’s classification method using their publicly available GitHub code, but used our own *control01* data as the training data. After obtaining predictions of expressed and unexpressed genes, we converted the classification results into binary expression values for our ‘Detection of subtype-specific signature genes’ analysis, assigning a value of 1 to predicted expressed genes and 0 to predicted unexpressed genes.

### Consideration of GC content as an input feature

Inclusion of GC content as an input feature into the SVR resulted in only negligible improvements in performance, and was, therefore, not included in final model.

### Consideration of LuCaP PDX inclusion in training

The current model was trained exclusively on control cfDNA, which may limit its ability to capture features derived from tumor cfDNA. To address this, we incorporated the LuCaP PDX samples (one CRPC-AR and one CRPC-NE sample with high tumor fraction)^26^ described above into the training dataset, together with two control samples^17^. This modification, however, also led to negligible improvements in performance based on similar Spearman correlation and MSE results (Fig. S4), and the same significance results of detecting subtype-specific signature genes (Fig. 5). However, this approach significantly reduces the applicability of our method to other cancer types, without the availability of cancer-specific PDX data for training. For example, pathway enrichment analysis on metastatic breast cancer (MBC) data resulted in enrichment of CRPC-specific pathways (i.e., BELTRAN_NEPC_UP), alongside relevant ones like SMID_BREAST_CANCER_LUMINAL_A_UP (Fig. S7d), indicating cancer type overfitting. Therefore, we chose not to incorporate these samples in training the final model.

### Pathway enrichment analysis using GSEA

GSEA was conducted to assess pathway activity in all cohorts: a CRPC-Adeno cohort, a CRPC-NE cohort, and an MBC cohort. cfOncoXpress gene expression predictions in each cohort were compared to the gene expression from RNA-seq of PBMCs from a panel of healthy controls (n = 34). Data were standardized using Min-Max normalization. GSEA (v.4.3.3)^50^ was applied with the following parameters: “No-Collapse”, Permutation_type = “gene_set”, Enrichment statistic = “weighted”, Metric for ranking genes = “Diff_of_Classes”, Max size = 500, and Min size = 1. Significant enrichment was defined as NOM p-value ≤ 0.05, FWER p-val ≤ 0.05, and FDR q-value ≤ 0.05.

We evaluated the enrichment scores for the Molecular Signatures Database (MSigDB) HALLMARK gene sets^55^. To capture key molecular features of CRPC, we included pathways associated with two major CRPC subtypes: the TCGA_ANDROGEN_RESPONSE pathway which is upregulated in CRPC-AR, and the BELTRAN_NEPC_UP pathway which is upregulated in CRPC-NE. These pathways were chosen to reflect the distinct transcriptional programs underlying CRPC heterogeneity and progression. Additionally, we selected five breast cancer-related pathways from the MSigDB C2 curated gene sets representing the luminal (SMID_BREAST_CANCER_LUMINAL_A_UP, SMID_BREAST_CANCER_LUMINAL_B_UP, CHARAFE_BREAST_CANCER_LUMINAL_VS_BASAL_UP), basal (HUPER_BREAST_BASAL_VS_LUMINAL_UP), and HER2-enriched (BIOCARTA_HER2_PATHWAY) subtypes. These pathways were chosen because they reflect the principal molecular subtypes of breast cancer, each with distinct biological behaviors, treatment responses, and clinical outcomes.

### Robustness and scalability of cfOncoXpress

#### Evaluating sequencing depth and tumor fraction requirements

To assess the sensitivity of our prediction correlation with respect to changes in tumor fraction and coverage, we performed in silico mixing of DTB-214-Baseline (125.33X, TFx = 0.6349) with a no-cancer control sample control01 (192.03X, TFx = 0) across various target coverage and tumor fraction combinations. The target tumor fractions evaluated were 0.05, 0.1, 0.2, 0.3, 0.4, 0.5, and 0.6. Target sequencing depths were generated as proportional fractions of the total patient sample coverage 125.33X), corresponding synthetic ultra-low-pass (ULP)WGS at approximately 0.251X coverage, low coverage (LC)_WGS at around 1.253X, intermediate coverage (6.266X to 25.066X), and standard WGS) at greater than 30X.

We compute the required fractions of DTB-214-Baseline and the control sample for mixing in order to achieve the desired target coverage and tumor fraction as follows: We define two samples—patient sample *S*_1_ with tumor fraction *T*_1_ and coverage *C*_1_, and healthy sample *S*_2_ with *T*_2_ =0 and coverage *C*_2_ . Our target mixed sample is *S*_3_, such that 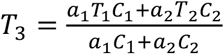 and *C*_3_ = *a*_1_*C*_1_ + *a*_2_*C*_2_, for some constants *a*_1_ and *a*_2_. This relies on the assumption that every sample is made up of some tumor and healthy component such that 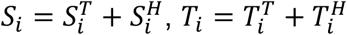, and 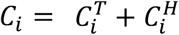. Then 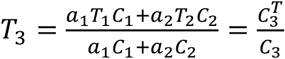. And *a*_1_*C*_1_ + *a*_2_*C*_2_ = *C*_3_, *a*_1_*T*_3_*C*_1_ + *a*_2_*T*_3_*C*_2_ = *a*_1_*T*_1_*C*_1_ + *a*_2_*T*_2_*C*_2_, and *C*_1_(*T*_3_ − *T*_1_)*a*_1_ + *C*_2_(*T*_3_ − *T*_2_)*a*_2_ = 0. Thus, we solve the following linear equation to solve for fractions *a*_1_ and 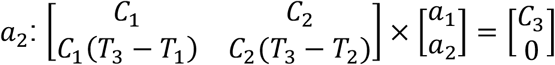.

#### Patient cohort size needed to identify differentially expressed genes

Power calculations were performed to estimate the patient sample size needed for pathway activity analysis given a panel of PBMC RNA-seq from a panel of healthy controls. That is, we estimate the required number of patient samples needed to identify differentially expressed genes in tumor samples when comparing against this PBMC panel. To do this calculation, we consider a dataset where samples are categorized into two groups: tumor and control, based on differential gene expression. Suppose each cluster can be modeled as a Gaussian distribution with mean (μ) and standard deviation (σ). The CRPC-Adeno^27^ was used to model the tumor data. The mean (μ_control_) and standard deviation (σ_control_) were derived from the expression values of genes in the TCGA_ANDROGEN_RESPONSE pathway in the PBMC panel. Thus, we calculate a series of μ_tumor_ values by simulating the fold change of the mean predicted expression of the TCGA_ANDROGEN_RESPONSE pathway genes in the CRPC-Adeno cohort relative to that of the PBMC panel (i.e., μ_tumor_ = foldchange * μ_control_). The standard deviation for the tumor cohort (σ_tumor_) was similarly derived from the CRPC-Adeno cohort. Using μ_control_, σ_control_, μ_tumor_, and σ_tumor_, we modeled the probabilities of correct classifications and misclassifications of samples as tumor versus control. Based on these probabilities and the known cohort size of healthy controls, we then estimated the required number of tumor samples to successfully distinguish between the two distributions (within set type I and type II errors). Thus, we obtain the required number of tumor samples as a function of fold-change (Fig. 8). When the fold-change of the mean predicted expression in tumor samples compared to controls is low (i.e. μ_tumor_ is close to μ_control_), more samples are required to distinguish between the distributions with confidence. Conversely, when the fold-change is high (i.e. μ_tumor_ is much higher from μ_control_), fewer samples are required to make this distinction (Fig. 8).

**Figure 8.**
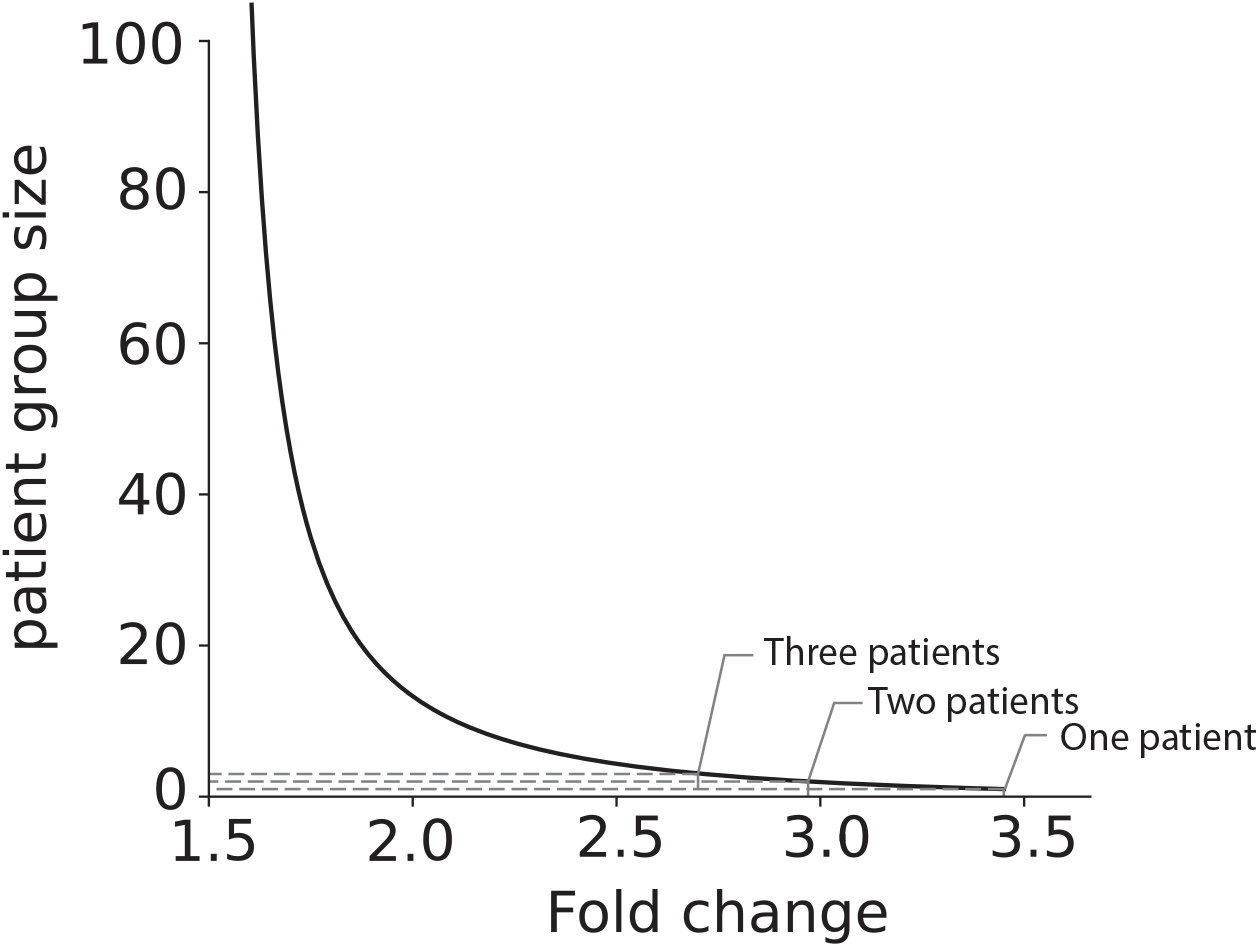
Estimated patient cohort sizes required to detect enriched pathways. It illustrates the minimum number of patients needed to detect differential pathway expression based on the fold change in predicted mean log_2_(expression) relative to log_2_(TPM) in healthy controls. Three reference points are as follows: a fold change of 3.45 requires one patient; 2.97 requires two patients; and 2.70 requires three patients. For example, CRPC-AR (TCGA_ANDROGEN_RESPONSE), CRPC-NE (BELTRAN_NEPC_UP), and Luminal A (SMID_BREAST_CANCER_LUMINAL_A_UP) pathways showed fold changes of 2.83, 5.08, and 3.35, respectively—requiring a minimum of three, one, and two patients for detection.

## Code availability

Relevant code is available at https://github.com/khuranalab/cfOncoXpress.

### Acknowledgements

We would like to thank Shahd ElNaggar, a summer student in Dr. Khurana’s lab, for downloading the cfDNA WGS data from CRPC cohort; and Hao Xu, for aligning the data to hg38. We also want to express our gratitude to Sandra Cohen, our wet lab manager, for her invaluable support.

## References

1. Kustanovich, A., Schwartz, R., Peretz, T. & Grinshpun, A. Life and death of circulating cell-free DNA. Cancer Biol Ther 20, 1057–1067 (2019).

2. Heitzer, E., Auinger, L. & Speicher, M. R. Cell-Free DNA and Apoptosis: How Dead Cells Inform About the Living. Trends Mol Med 26, 519–528 (2020).

3. Alix-Panabières, C., Schwarzenbach, H. & Pantel, K. Circulating Tumor Cells and Circulating Tumor DNA. Annu Rev Med 63, 199–215 (2012).

4. Haber, D. A. & Velculescu, V. E. Blood-Based Analyses of Cancer: Circulating Tumor Cells and Circulating Tumor DNA. Cancer Discov 4, 650–661 (2014).

5. Bettegowda, C. et al. Detection of Circulating Tumor DNA in Early- and Late-Stage Human Malignancies. Sci Transl Med 6, (2014).

6. Heitzer, E., Ulz, P. & Geigl, J. B. Circulating Tumor DNA as a Liquid Biopsy for Cancer. Clin Chem 61, 112–123 (2015).

7. Alix-Panabières, C. & Pantel, K. Clinical Applications of Circulating Tumor Cells and Circulating Tumor DNA as Liquid Biopsy. Cancer Discov 6, 479–491 (2016).

8. Corcoran, R. B. & Chabner, B. A. Application of Cell-free DNA Analysis to Cancer Treatment. New England Journal of Medicine 379, 1754–1765 (2018).

9. Bronkhorst, A. J., Ungerer, V. & Holdenrieder, S. The emerging role of cell-free DNA as a molecular marker for cancer management. Biomol Detect Quantif 17, 100087 (2019).

10. Lone, S. N. et al. Liquid biopsy: a step closer to transform diagnosis, prognosis and future of cancer treatments. Mol Cancer 21, 79 (2022).

11. Lo, Y. M. D. et al. Maternal Plasma DNA Sequencing Reveals the Genome-Wide Genetic and Mutational Profile of the Fetus. Sci Transl Med 2, (2010).

12. Snyder, M. W., Kircher, M., Hill, A. J., Daza, R. M. & Shendure, J. Cell-free DNA Comprises an In Vivo Nucleosome Footprint that Informs Its Tissues-Of-Origin. Cell 164, 57–68 (2016).

13. Schones, D. E. et al. Dynamic Regulation of Nucleosome Positioning in the Human Genome. Cell 132, 887–898 (2008).

14. Valouev, A. et al. Determinants of nucleosome organization in primary human cells. Nature 474, 516–520 (2011).

15. Venkatesh, S. & Workman, J. L. Histone exchange, chromatin structure and the regulation of transcription. Nat Rev Mol Cell Biol 16, 178–189 (2015).

16. Ulz, P. et al. Inferring expressed genes by whole-genome sequencing of plasma DNA. Nat Genet 48, 1273–1278 (2016).

17. Esfahani, M. S. et al. Inferring gene expression from cell-free DNA fragmentation profiles. Nat Biotechnol 40, 585–597 (2022).

18. Henrichsen, C. N., Chaignat, E. & Reymond, A. Copy number variants, diseases and gene expression. Hum Mol Genet 18, R1–R8 (2009).

19. Gamazon, E. R. & Stranger, B. E. The impact of human copy number variation on gene expression. Brief Funct Genomics 14, 352–7 (2015).

20. Shao, X. et al. Copy number variation is highly correlated with differential gene expression: a pancancer study. BMC Med Genet 20, 175 (2019).

21. Bailey, C. et al. Origins and impact of extrachromosomal DNA. Nature 635, 193–200 (2024).

22. Turner, K. M. et al. Extrachromosomal oncogene amplification drives tumour evolution and genetic heterogeneity. Nature 543, 122–125 (2017).

23. Kim, H. et al. Extrachromosomal DNA is associated with oncogene amplification and poor outcome across multiple cancers. Nat Genet 52, 891–897 (2020).

24. Wu, S. et al. Circular ecDNA promotes accessible chromatin and high oncogene expression. Nature 575, 699–703 (2019).

25. Lange, J. T. et al. The evolutionary dynamics of extrachromosomal DNA in human cancers. Nat Genet 54, 1527–1533 (2022).

26. De Sarkar, N. et al. Nucleosome Patterns in Circulating Tumor DNA Reveal Transcriptional Regulation of Advanced Prostate Cancer Phenotypes. Cancer Discov 13, 632–653 (2023).

27. Herberts, C. et al. Deep whole-genome ctDNA chronology of treatment-resistant prostate cancer. Nature 608, 199–208 (2022).

28. Zivanovic Bujak, A. et al. Circulating tumour DNA in metastatic breast cancer to guide clinical trial enrolment and precision oncology: A cohort study. PLoS Med 17, e1003363 (2020).

29. Adalsteinsson, V. A. et al. Scalable whole-exome sequencing of cell-free DNA reveals high concordance with metastatic tumors. Nat Commun 8, 1324 (2017).

30. Cristiano, S. et al. Genome-wide cell-free DNA fragmentation in patients with cancer. Nature 570, 385–389 (2019).

31. Akiba, T., Sano, S., Yanase, T., Ohta, T. & Koyama, M. Optuna. in Proceedings of the 25th ACM SIGKDD International Conference on Knowledge Discovery & Data Mining 2623–2631 (ACM, 2019). doi:10.1145/3292500.3330701.

32. Talevich, E., Shain, A. H., Botton, T. & Bastian, B. C. CNVkit: Genome-Wide Copy Number Detection and Visualization from Targeted DNA Sequencing. PLoS Comput Biol 12, e1004873 (2016).

33. Luebeck, J. et al. AmpliconSuite: an end-to-end workflow for analyzing focal amplifications in cancer genomes. bioRxiv 2024.05.06.592768 (2024) doi:10.1101/2024.05.06.592768.

34. Zhao, S. G. et al. Integrated analyses highlight interactions between the three-dimensional genome and DNA, RNA and epigenomic alterations in metastatic prostate cancer. Nat Genet 56, 1689– 1700 (2024).

35. Storlazzi, C. T. et al. Gene amplification as double minutes or homogeneously staining regions in solid tumors: origin and structure. Genome Res 20, 1198–206 (2010).

36. Campbell, P. J. et al. Identification of somatically acquired rearrangements in cancer using genome-wide massively parallel paired-end sequencing. Nat Genet 40, 722–9 (2008).

37. Bignell, G. R. et al. Architectures of somatic genomic rearrangement in human cancer amplicons at sequence-level resolution. Genome Res 17, 1296–303 (2007).

38. Li, Z., Wang, B., Liang, H. & Han, L. Pioneering insights of extrachromosomal DNA (ecDNA) generation, action and its implications for cancer therapy. Int J Biol Sci 18, 4006–4025 (2022).

39. Qiu, X. et al. MYC drives aggressive prostate cancer by disrupting transcriptional pause release at androgen receptor targets. Nat Commun 13, 2559 (2022).

40. Djusberg, E. et al. High levels of the AR-V7 Splice Variant and Co-Amplification of the Golgi Protein Coding YIPF6 in AR Amplified Prostate Cancer Bone Metastases. Prostate 77, 625–638 (2017).

41. Kakarla, M. et al. Ephrin B Activate Src Family Kinases in Fibroblasts Inducing Stromal Remodeling in Prostate Cancer. Cancers (Basel) 14, (2022).

42. Liu, J. et al. Androgen deprivation-induced OPHN1 amplification promotes castration-resistant prostate cancer. Oncol Rep 47, (2022).

43. Archer, M. et al. Kinesin Facilitates Phenotypic Targeting of Therapeutic Resistance in Advanced Prostate Cancer. Mol Cancer Res 22, 730–745 (2024).

44. Pornour, M. et al. USP11 promotes prostate cancer progression by up-regulating AR and c-Myc activity. Proc Natl Acad Sci U S A 121, e2403331121 (2024).

45. Das, R. et al. An integrated functional and clinical genomics approach reveals genes driving aggressive metastatic prostate cancer. Nat Commun 12, 4601 (2021).

46. Chen, L.-J. et al. The role of lysine-specific demethylase 6A (KDM6A) in tumorigenesis and its therapeutic potentials in cancer therapy. Bioorg Chem 133, 106409 (2023).

47. Tang, F. et al. Chromatin profiles classify castration-resistant prostate cancers suggesting therapeutic targets. Science (1979) 376, (2022).

48. Esfahani, M. S. et al. Inferring gene expression from cell-free DNA fragmentation profiles. Nat Biotechnol 40, 585–597 (2022).

49. Ulz, P. et al. Inferring expressed genes by whole-genome sequencing of plasma DNA. Nat Genet 48, 1273–1278 (2016).

50. Subramanian, A. et al. Gene set enrichment analysis: A knowledge-based approach for interpreting genome-wide expression profiles. Proceedings of the National Academy of Sciences 102, 15545–15550 (2005).

51. Monaco, G. et al. RNA-Seq Signatures Normalized by mRNA Abundance Allow Absolute Deconvolution of Human Immune Cell Types. Cell Rep 26, 1627-1640.e7 (2019).

52. Zucca, S. et al. RNA-Seq profiling in peripheral blood mononuclear cells of amyotrophic lateral sclerosis patients and controls. Sci Data 6, 190006 (2019).

53. Han, Z. et al. RNA-seq profiling reveals PBMC RNA as a potential biomarker for hepatocellular carcinoma. Sci Rep 11, 17797 (2021).

54. Beltran, H. et al. Divergent clonal evolution of castration-resistant neuroendocrine prostate cancer. Nat Med 22, 298–305 (2016).

55. Liberzon, A. et al. The Molecular Signatures Database Hallmark Gene Set Collection. Cell Syst 1, 417–425 (2015).

56. Johnson, K. S., Conant, E. F. & Soo, M. S. Molecular Subtypes of Breast Cancer: A Review for Breast Radiologists. J Breast Imaging 3, 12–24 (2021).

57. Heintzman, N. D. et al. Distinct and predictive chromatin signatures of transcriptional promoters and enhancers in the human genome. Nat Genet 39, 311–318 (2007).

58. Jin, C. et al. H3.3/H2A.Z double variant–containing nucleosomes mark ‘nucleosome-free regions’ of active promoters and other regulatory regions. Nat Genet 41, 941–945 (2009).

59. Ong, C.-T. & Corces, V. G. Enhancer function: new insights into the regulation of tissue-specific gene expression. Nat Rev Genet 12, 283–293 (2011).

60. Andersson, R. & Sandelin, A. Determinants of enhancer and promoter activities of regulatory elements. Nat Rev Genet 21, 71–87 (2020).

61. Xu, D. et al. Recapitulation of patient-specific 3D chromatin conformation using machine learning. Cell Reports Methods 3, 100578 (2023).

62. Quigley, D. A. et al. Genomic Hallmarks and Structural Variation in Metastatic Prostate Cancer. Cell 174, 758-769.e9 (2018).

63. Sarkar, N. De et al. Nucleosome Patterns in Circulating Tumor DNA Reveal Transcriptional Regulation of Advanced Prostate Cancer Phenotypes. Cancer Discov 13, 632–653 (2023).

64. Akiba, T., Sano, S., Yanase, T., Ohta, T. & Koyama, M. Optuna. in Proceedings of the 25th ACM SIGKDD International Conference on Knowledge Discovery & Data Mining 2623–2631 (ACM, New York, NY, USA, 2019). doi:10.1145/3292500.3330701.

65. Abadi, M. et al. TensorFlow: Large-Scale Machine Learning on Heterogeneous Distributed Systems. (2016).

