## supplementary figures for "cfOncoXpress: Tumor gene expression prediction from cell-free DNA whole-genome sequences"

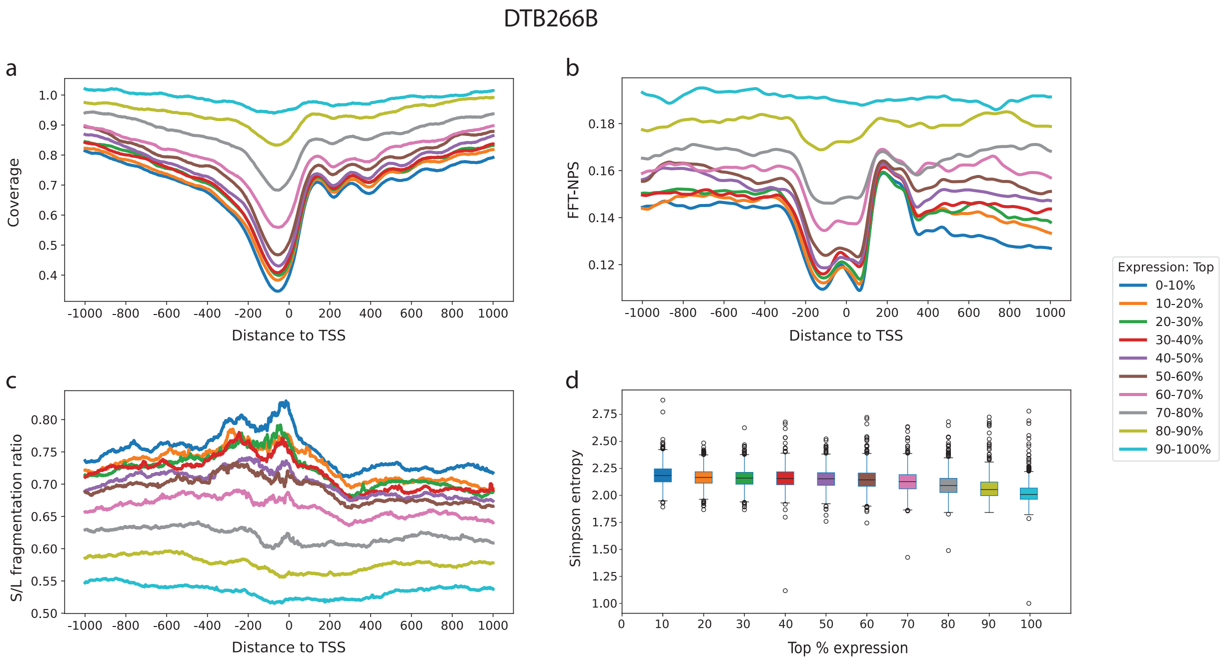


**Figure S1. cfDNA features of DTB266B (tumor fraction = 0.12) near transcription start sites correlate with gene expression.**

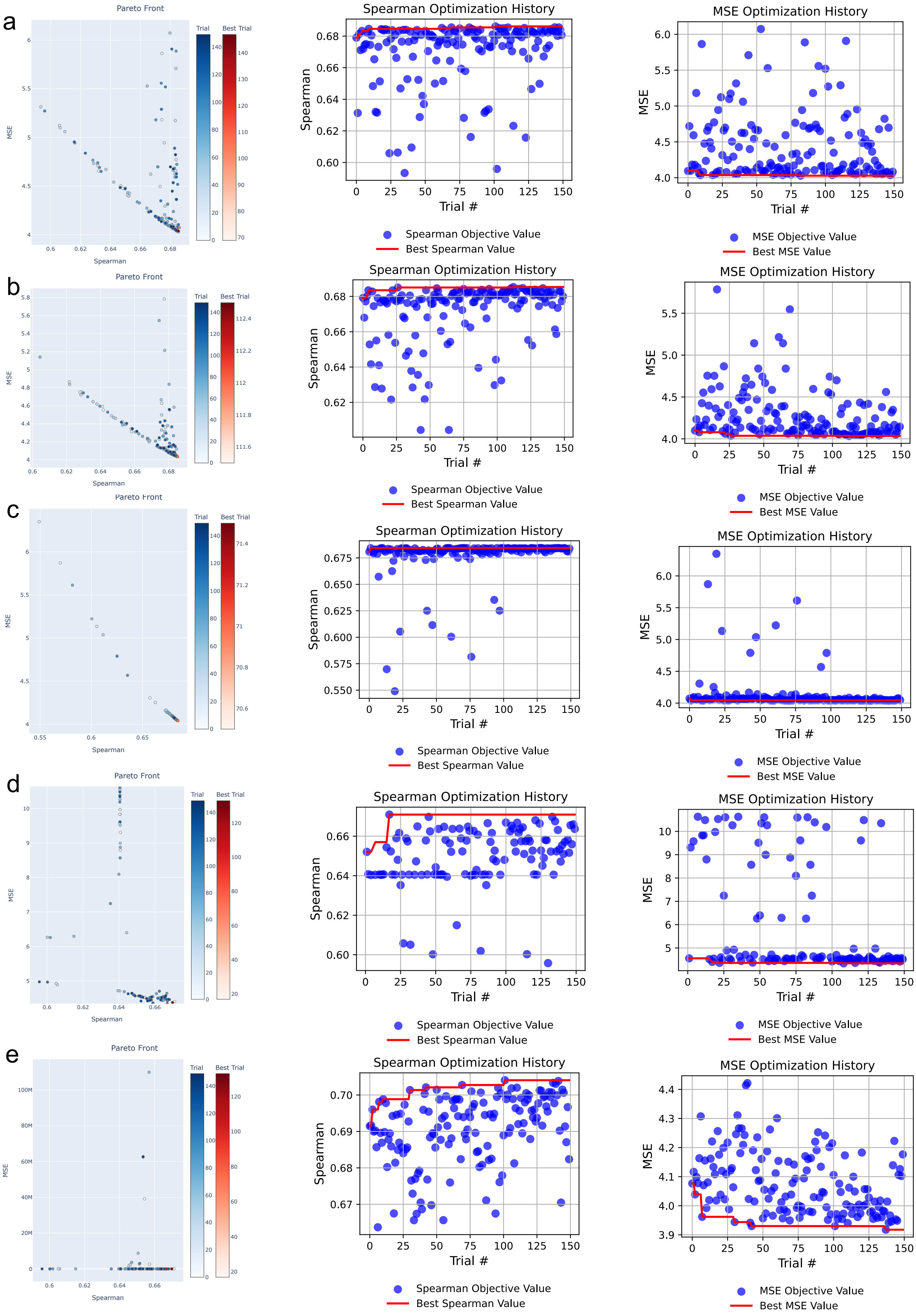

a
b
c
d
e
Pareto Front
Trial
Best Trial
MSE
Spearman
Spearman Optimization History
Spearman
Trial #
Spearman Objective Value
Best Spearman Value
MSE Optimization History
MSE
Trial #
MSE Objective Value
Best MSE Value

**Figure S2. Optuna hyperparameter optimization by checking the performance of spearman correlation and mean squared error (MSE) between predicted and observed gene expression values.** The best result for (a) gradient boosting model is spearman r = 0.686, MSE = 4.037 with the optimal params = {'n_estimators': 735, 'max_depth': 9, 'min_samples_split': 8, 'min_samples_leaf': 14, 'learning_rate': 0.00428, 'subsample': 0.608, 'max_features': 'log2'};
(b) extreme gradient boosting is spearman r = 0.685, MSE = 4.033 with the optimal params = {'n_estimators': 696, 'max_depth': 5, 'learning_rate': 0.0114, 'subsample': 0.775, 'colsample_bytree': 0.77, 'reg_alpha': 0.00262, 'reg_lambda': 0.0344, 'min_child_weight': 5};
(c) random forest is spearman r = 0.684, MSE = 4.042, with the optimal params = {'n_estimators': 824, 'max_depth': 11, 'min_samples_split': 7, 'min_samples_leaf': 5, 'max_features': 'sqrt', 'bootstrap': True};
(d) support vector regression is spearman r = 0.671, MSE = 4.404 with the optimal params = {'C': 397.1039921978707, 'epsilon': 0.7351783229677453, 'kernel': 'rbf', 'gamma': 'scale'};
(e) neuron network is spearman r = 0.704, MSE = 3.93 with the optimal params = {'n_layers': 3, 'units_l0': 80, 'act_l0': 'relu', 'dropout_l0': 0.157, 'units_l1': 96, 'act_l1': 'tanh', 'dropout_l1': 0.220, 'units_l2': 128, 'act_l2': 'tanh', 'dropout_l2': 0.257, 'learning_rate': 0.000353, 'batch_size': 32}.
Neuron network gets the best performance with highest spearman correlation with log2 TPM and lowest mean square errors in training dataset.

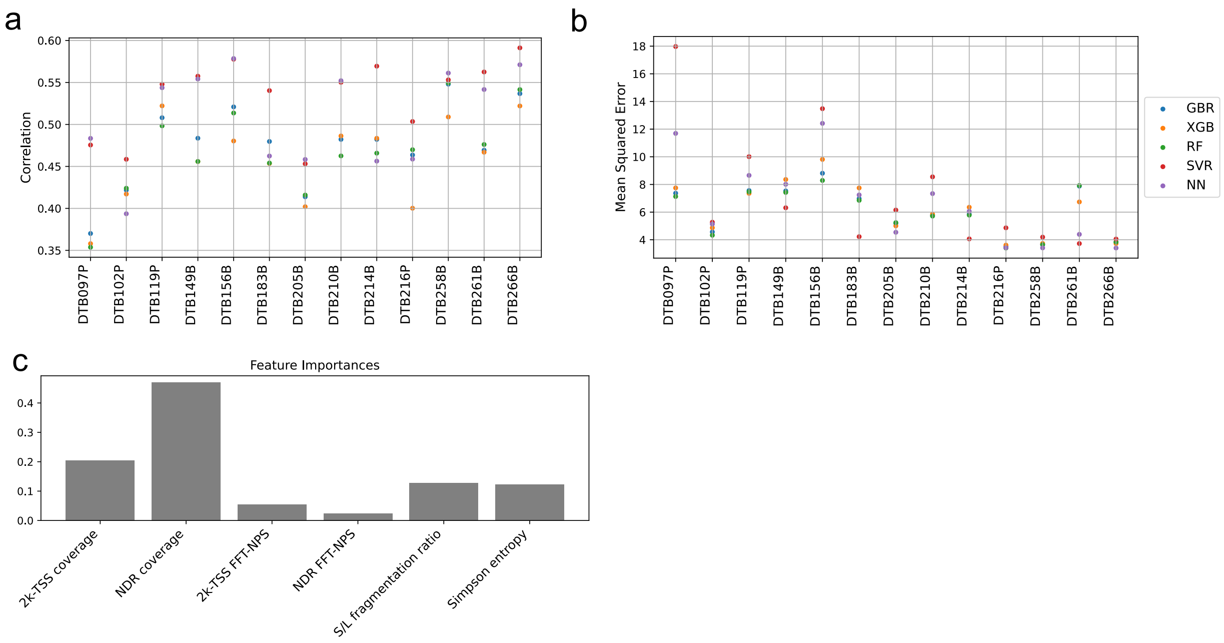


**Figure S3. Hyperparameter tuning and optimal model chosen.** Evaluation of model performance on the test set in terms of (a) spearman correlation and (b) mean squared error. (c) Permutation feature importance was used to evaluate the contribution of six cfDNA features, with the following ranking of importance: NDR coverage > 2k-TSS coverage > S/L fragmentation ratio > Simpson entropy > 2k-TSS FFT-NPS > NDR FFT-NPS.

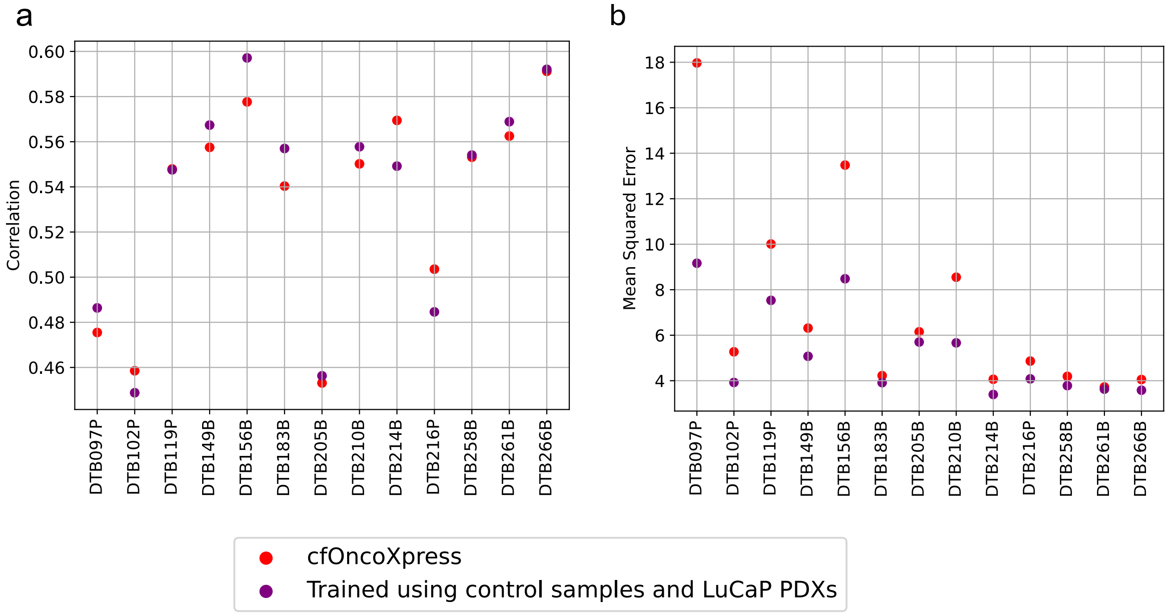


**Figure S4. Model performance was evaluated using Spearman correlation and mean squared error between predicted expression and $log_2$(TPM+1) from matched tissue RNA-seq.** Incorporating controls and LuCaP PDXs during training increased correlation by less than 0.02 and yielded a modest reduction in MSE, but overall performance did not improve substantially.

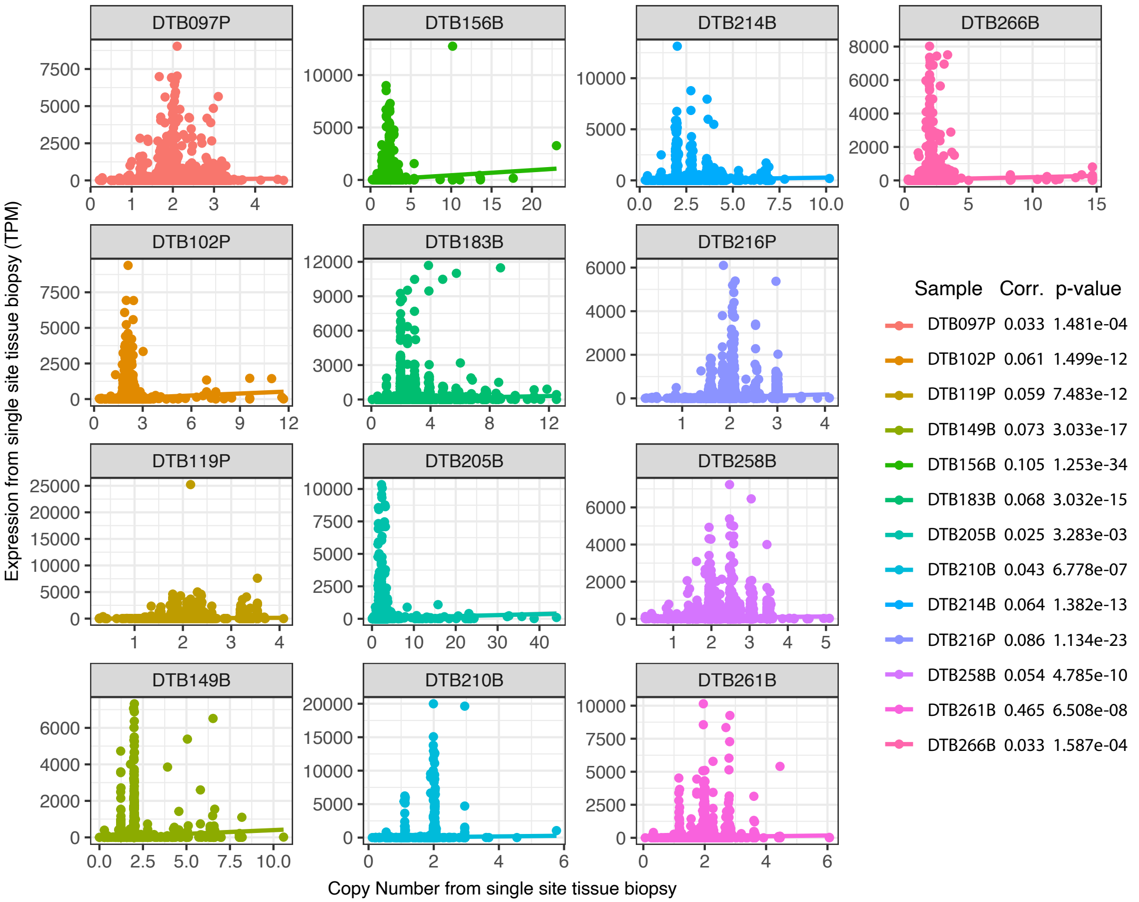


- **Figure S5.** There's no correlation (Pearson) between copy number from single-site tissue biopsy and tissue RNA-seq gene expression.

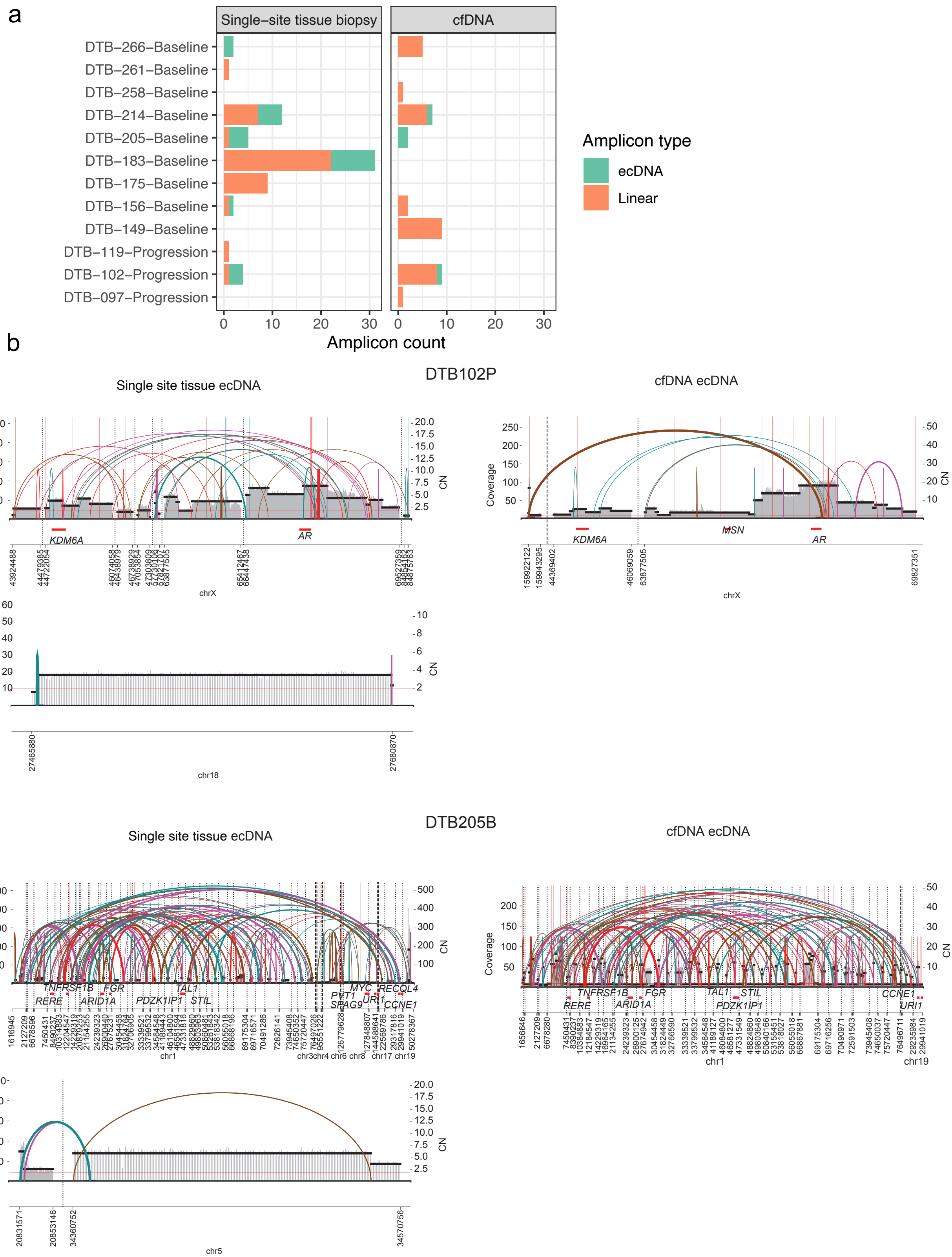
a
Single−site tissue biopsy
cfDNA
DTB−266−Baseline
DTB−261−Baseline
DTB−258−Baseline
DTB−214−Baseline
DTB−205−Baseline
DTB−183−Baseline
DTB−175−Baseline
DTB−156−Baseline
DTB−149−Baseline
DTB−119−Progression
DTB−102−Progression
DTB−097−Progression
0
10
20
30
Amplicon count
Amplicon type
ecDNA
Linear
b
DTB102P
Single site tissue ecDNA
cfDNA ecDNA
Coverage
CN
KDM6A
AR
MSN
chrX
chr18
27465880
27680870
DTB205B
Single site tissue ecDNA
cfDNA ecDNA
TNFRSF1B
RERE
FGR
ARID1A
TAL1
PDZK1IP1
STIL
PVT1
SPAG9
MYC
URI1
RECQL4
CCNE1
chr1
chr19
chr5
20831571
20853146
34360752
34570756

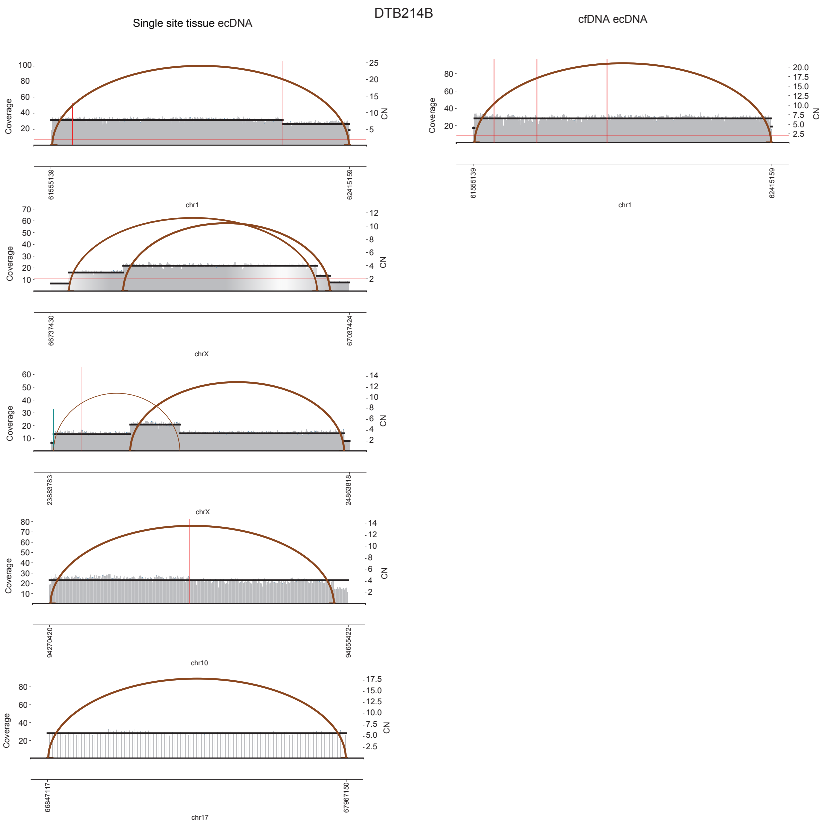


**Figure S6.** (a) Detected amplicons from tissue biopsy and cfDNA, ecDNA (green) and linear amplicons (orange). (b) 3 samples' ecdna detected from tissue vs. cfDNA.

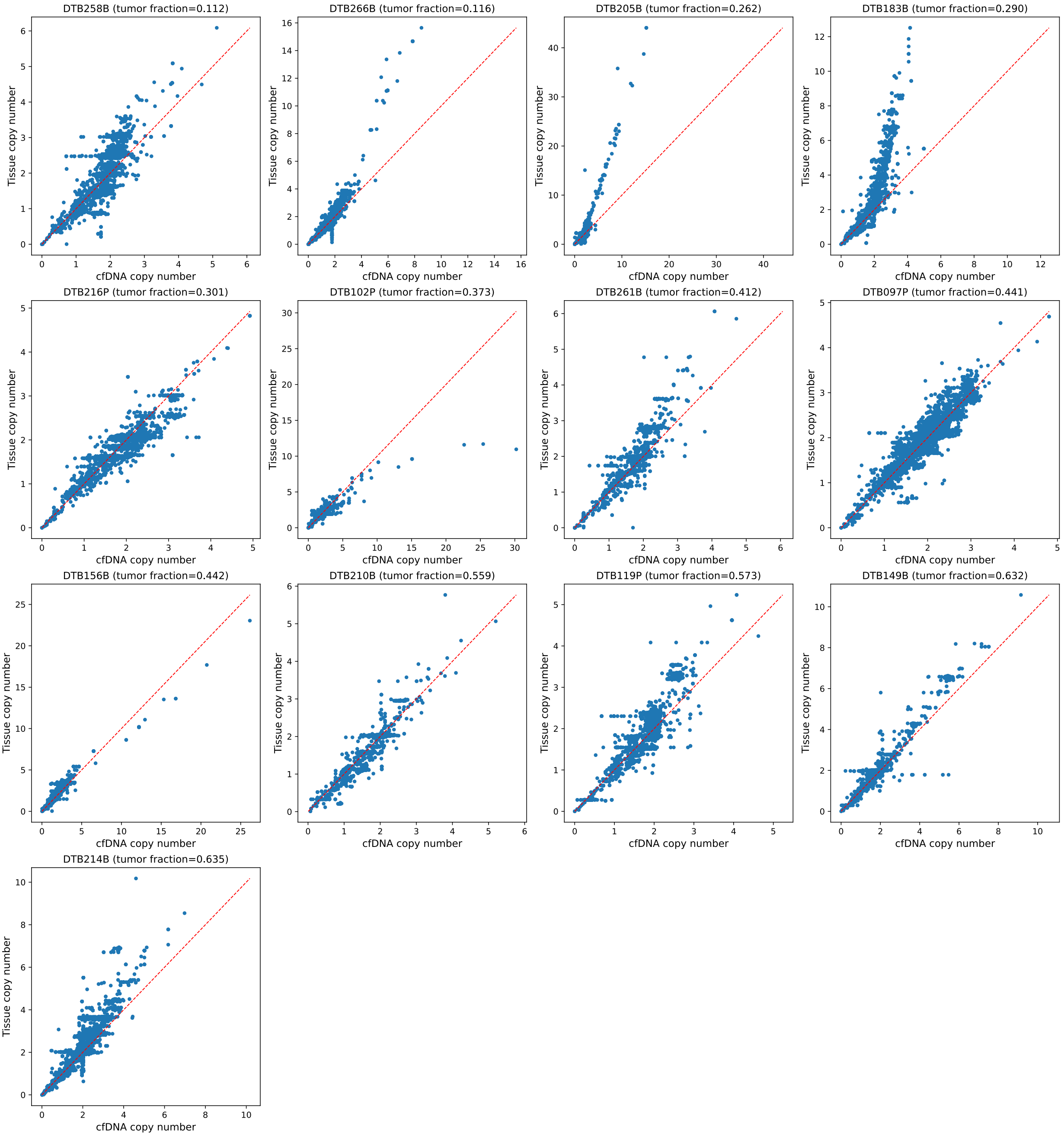


**Figure S7. Tissue copy number is linearly correlated with cfDNA copy number.** Pearson correlations between cfDNA and tissue copy number range from 0.803 to 0.954. Panels are arranged in order of increasing tumor fraction.
